# Sugar transporter Slc37a2 regulates bone metabolism via a dynamic tubular lysosomal network in osteoclasts

**DOI:** 10.1101/2022.04.28.489831

**Authors:** Pei Ying Ng, Amy B.P. Ribet, Qiang Guo, Benjamin H. Mullin, Jamie W.Y. Tan, Euphemie Landao-Bassonga, Sébastien Stephens, Kai Chen, Laila Abudulai, Maike Bollen, Edward T.T.T. Nguyen, Jasreen Kular, John M. Papadimitriou, Kent Søe, Rohan D. Teasdale, Jiake Xu, Robert G. Parton, Hiroshi Takanayagi, Nathan J. Pavlos

## Abstract

Osteoclasts are giant bone-digesting cells that harbour specialized lysosome-related organelles termed secretory lysosomes (SLs). SLs store cathepsin K and serve as a membrane precursor to the ruffled border, the osteoclast’s ‘resorptive apparatus’. Yet, the molecular composition and spatiotemporal organization of SLs remains incompletely understood. Here, using organelle-resolution proteomics, we identify member a2 of the solute carrier 37 family (Slc37a2) as a SL sugar transporter. We demonstrate that Slc37a2 localizes to the SL limiting membrane and that these organelles adopt a hitherto unnoticed but dynamic tubular network in living osteoclasts that is required for bone digestion. Accordingly, mice lacking Slc37a2 accrue high bone mass owing to uncoupled bone metabolism and disturbances in SL export of monosaccharide sugars, a prerequisite for SL delivery to the ruffled border. Thus, Slc37a2 is a physiological component of the osteoclast’s unique secretory organelle and a potential therapeutic target for metabolic bone diseases.

## INTRODUCTION

Osteoclasts are bone-digesting cells that play a central role in skeletal bone growth and metabolism and underscore pathologies such as osteoporosis and osteopetrosis (Zaidi, 2007, Teitelbaum and Ross, 2003, Cappariello et al., 2014). These multinucleated giants degrade bone via the ruffled border, a villous-like plasma membrane domain circumscribed by an actin ring that serves as the osteoclast’s unique ‘resorptive apparatus’. Ruffled border formation involves the polarized fusion of specialized lysosomal-related organelles (LROs), termed secretory lysosomes (SLs), with the bone-apposed plasmalemma (Stenbeck, 2002, Baron et al., 1988). Unlike conventional lysosomes, which traditionally serve as intracellular depots for the degradation and recycling of endogenous and exogenous biomolecules (Saftig and Klumperman, 2009), LROs are ‘hybrid organelles’ that share features of both late-endosomes and lysosomes coupled with exocytic functions (Delevoye et al., 2019). Osteoclast SLs store the acidic hydrolase cathepsin K (Drake et al., 1996) and are compositionally and functionally defined by the presence of several endo-lysosomal membrane proteins, namely LAMP1/2 (Palokangas et al., 1997), Rab7 (Zhao et al., 2001), the *a3* V-ATPase proton pump subunit (Toyomura et al., 2003), chloride channel ClC-7 (Kornak et al., 2001), TI-VAMP/VAMP7 and synaptotagmin 7 (Zhao et al., 2008). Fusion of SLs with the bone-apposed plasmalemma discharges cathepsin K into the underlying resorptive microenvironment to digest collagenous (Type 1a) bone matrix. At the same time, coalescence of SL membranes with the ventral plasmalemma enriches the ruffled border with nanoscale bone-digesting machinery, including V-ATPases (Blair et al., 1989) and ClC-7 (Kornak et al., 2001), which cooperatively acidify the resorptive space to dissolve bone mineral. The ruffled border, together with the juxtaposed resorption lacuna, is therefore considered a giant digestive ‘extracellular LRO’ (Baron et al., 1985).

As a precursor membrane for ruffled border genesis, SLs represent fertile grounds for the discovery of new homeostatic regulators of bone mass and thus potential anti-resorptive drug targets. Yet despite their critical importance, our understanding of osteoclast SLs remains limited. In particular, compared to other well-established LROs (e.g. melanosomes in melanocytes) (Delevoye et al., 2019), we still lack elementary information regarding SL composition, organization and regulation. Thus far, the molecular anatomy of SLs has largely been gleamed from unsystematic studies of genetically-modified mice with high bone mass phenotypes and incidental genetic findings arising from patients with rare sclerosing bone dysplasia (Sobacchi et al., 2013). While these studies have greatly advanced our understanding of the molecules involved in SL biogenesis (Lacombe et al., 2013) as well as the participants and downstream effectors regulating SL trafficking (Ng et al., 2019), surprisingly little is known about the nature and number of transport systems that reside on SLs and orchestrate the exchange of small molecules, such as ions, amino acids and sugars across its membrane (Ribet et al., 2021). In addition, how SLs are organized in osteoclasts, both spatially and temporally, is unclear.

Here, to expand the molecular landscape of osteoclast SLs, we systematically surveyed the proteome of enriched SLs isolated from murine osteoclasts and identified Slc37a2 as a candidate SL transporter. Unexpectedly, using live cell imaging, we reveal that Slc37a2^+^ SLs exist as a hitherto unreported but highly dynamic tubular lysosomal network in osteoclasts that radiates throughout the cytoplasm and fuses with the bone-apposed plasmalemma. Moreover, deletion of *Slc37a2* in mice results in a profound high bone mass phenotype attributed to osteoclast dysfunction and associated disturbances in SL resolution and delivery to the ruffled border. Altogether, our findings unmask Slc37a2 as a SL sugar transporter critical for bone metabolism and highlight previously unappreciated plasticity of the osteoclast’s specialized lysosome-related organelle(s).

## RESULTS

### Proteomics of osteoclast SLs identifies Slc37a2 as a candidate regulator of bone mass

To survey the proteome of osteoclast SLs, we adapted two well-established lysosome enrichment protocols using <u>s</u>uper<u>p</u>aramagnetic iron <u>o</u>xide <u>n</u>anoparticles (SPIONs) (Nakamura et al., 2014, Walker and Lloyd-Evans, 2015). For this, large scale murine bone marrow monocyte (BMM)-derived pre-osteoclast cultures (Day-3, post-RANKL stimulation) were ‘pulsed’ for 24 hours with SPIONs to encourage uptake into endosomes. As a derivative of endomembranes, we hypothesized that endocytosed nanoparticles could be ‘chased’ into SLs upon the convergence of SPION-loaded endosomes with lysosomes and secretory pathways, a process that is synchronized with the final stages of RANKL-mediated differentiation of pre-osteoclasts into mature αvβ_3_-integrin-positive (IntegriSense^645^) osteoclasts (>90% of cells) (**Fig. 1A**). Following a 24 hour ‘chase’ (total ‘pulse-chase’=48 hours), osteoclasts were homogenized, SPION-loaded organelles captured-from post-nuclear supernatants (PNS) using magnetic columns, and membranes serially eluted in fractions (F1-F3). The enrichment of isolated SPIONS-loaded organelles in the Fraction F2 preparation was confirmed by transmission electron microscopy (TEM) and their origins assessed immunoanalytically (**Fig. 1B**). Gratifyingly, captured F2-organelles showed substantial enrichment with major SL membrane markers including LAMP2 and Rab7, as well as the primary osteoclast luminal acidic hydrolase cathepsin K (Ctsk). This fraction was also highly enriched for the LRO resident GTPase Rab38 (Marks, 2012) but virtually devoid of contaminating organelles including early-endosomes (Rab5), ER-Golgi (Rab1b), Golgi (Syntaxin 16b), mitochondria (VDAC3), cytoskeleton (β-actin) and plasma membrane (V-Glut-1), thus deeming it suitable for proteomics.

**Fig. 1.**
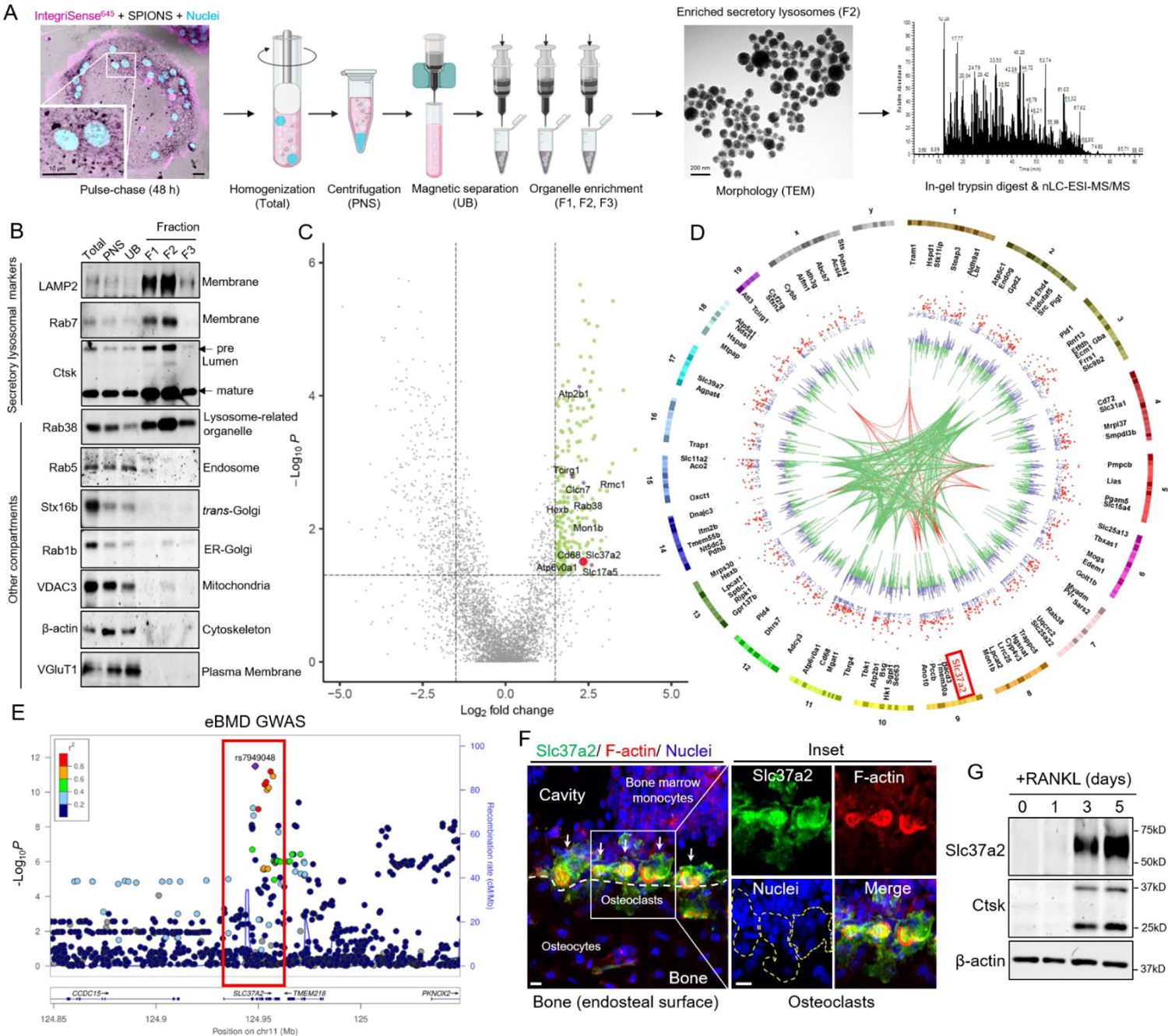
Proteomic analysis of enriched osteoclast SLs identifies Slc37a2 as a candidate regulator of bone mass. (A) Schematic of the workflow for the SPION-based SL enrichment method, validation and proteomic analysis. (B) Immunoblotting for protein markers of various subcellular compartments in whole-cell homogenates (H), post-nuclear supernatant (PNS), magnetic column flow-through (UB) and enriched SL fractions (F1, F2, F3). 5 µg of proteins of each fraction was analyzed by SDS-PAGE and immunoblotted with indicated antibodies. (C) Volcano plot depicting the difference between proteins in whole cell homogenates and SLs (*n* = 3). The top 218 proteins up-regulated in the lysosomal fractions are shown in green (FC >1.5, *P*<0.05) with top 6 membrane transporters colored in purple and Slc37a2 is highlighted in red. (D) CIRCOS plot (Krzywinski et al., 2009) displaying mouse chromosomes, gene names for the top 218 proteins up-regulated in the SL fractions, scatterplot representing protein abundance ratio (SLs/homogenate) adjusted *P* values (purple) with significant associations (*P* <0.05) colored red, histogram representing Log_2_FC abundance values for each protein SLs/homogenate purple (+ve), green (-ve), gene groupings representing human homologues associated with eBMD at *P*<6.6E^-9^ (green) and *P*<5E^-20^ (red) (Morris et al., 2019). Protein abundance ratio adjusted *P* values are displayed as -Log_10_ *P* values. (E) Regional association plots for the human *SLC37A2* gene (red box) generated using eBMD GWAS association results from (Morris et al., 2019). Genetic variants within 100kb of the lead variant (rs7949048, purple) are depicted (x axis) along with their eBMD *P* value (–Log_10_). (F) Slc37a2 expression in femur of 12-week old male WT mice by fluorescence immunohistochemistry. Arrows indicate osteoclasts and a dashed line defines bone (white) and osteoclast (yellow) surfaces. Scale bars, 10 μm. (G) Slc37a2 protein expression during *in vitro* osteoclast differentiation of primary BMMs by immunoblotting. Cathepsin K (Ctsk) and β-actin served as controls.

The protein composition of SL-enriched F2 fractions (from three biological replicates) was analyzed by high-resolution mass spectrometry (nLC-ESI-MS/MS). In total, 4153 different proteins were unambiguously identified defined by at least two unique peptides and master proteins. Of these, 218 achieved >1.5 Log_2_fold change (FC) and adjusted *P*<0.05 relative enrichment in SL F2-fraction compared to the starting cell homogenate by label-free quantitation (**Fig. 1C**). The results are summarized in **Extended Data Table 1**. As expected, the SL proteome was enriched with membrane proteins, including major SL resident proteins CIC-7 (Clcn7, enriched 2.35 Log_2_FC, *P*=3.51E^-02^) and V-ATPase 116 kDa *a3* subunit (Tcirg1, enriched 2.01 Log_2_FC, *P* =1.66E^-03^), thus validating our approach. In addition, the SL proteome was enriched with proteins involved in trafficking (n=14) including Rab38 (enriched 2.62 Log_2_FC, *P*=3.56E^-03^) as well as core subunits of the mammalian guanidine exchange factor for Rab7 i.e. Mon1b (enriched 2.66 Log_2_FC, *P*=7.63E^-03^) (Nordmann et al., 2010) and Regulator of Mon1-Ccz1 complex (Rmc1/*C18orf8;* enriched 3.06 Log_2_FC, *P*=2.72E^-03^) (van den Boomen et al., 2020) (**Fig. 1C**). Intriguingly, the SL proteome was dominated by proteins involved in molecular transport (n=43), with almost half representing members of the solute carrier (Slc) protein superfamily of secondary active transporters (n=16) (Pizzagalli et al., 2021) **(Extended Data Table 1)**. Finally, several miscellaneous proteins were also identified, some of which constitute major cargo of endolysosomes, for example, β-hexosaminidase (enriched 1.74 Log_2_FC, *P*=4.05E^-03^) and others that are predicted to localize to endolysosomal membranes but whose SL residency remains to be confirmed (e.g. CD68, enriched 2.25 Log_2_FC, *P*=1.95E^-02^) (**Fig. 1C**).

To first gauge the relevance of the mouse osteoclast SL proteome to humans, we investigated whether significant associations identified in a large human genome-wide association study (GWAS) for estimated bone mineral density (eBMD) (Morris et al., 2019) were enriched among human homologues of the mouse osteoclast SL proteome gene set. The top 218 proteins enriched in the SL F2-fraction are mapped onto a CIRCOS plot, with gene groupings representing those with human homologues strongly associated (*P*< 6.6E^-9^, green) and very strongly associated (*P*< 5E^-20^, red) with estimated BMD indicated (**Fig. 1D**). Using the VEGAS2Pathway approach (Mishra and MacGregor, 2017), we confirmed statistically significant enrichment of eBMD association signals in the SL gene set (*P EMPIRICAL*=0.04), indicating that the SL proteome is conserved in humans and that it is enriched with candidates clinically-relevant to the regulation of human bone mass.

To further validate the SL proteome, we next focused our attention on membrane transport proteins. We chose this protein family for two reasons: (i) they are polytopic proteins that contain multiple transmembrane domains (TMDs) and are thus integral components of the SL limiting membrane and; (ii) mutations in genes encoding SL transporters underscore most (>70%) of the known forms of human autosomal recessive osteopetrosis (ARO) (Sobacchi et al., 2013). To this end, the SL proteome was further filtered for ‘Membrane Transport Proteins’ that: (i) possessed >1 TMD (n=42); ii) housed a lysosomal targeting signal(s) outside of their TMDs (i.e. YxxØ or [DE]xxxL[LI], where Ø stands for an amino acid residue with a bulky hydrophobic side chain) (n=22); and; (iii) whose corresponding lead SNPs reached strong genome-wide significance in the human eBMD dataset (*P*<6.6E^-9^). Six candidates satisfied these criteria (**Fig. 1C-D**): (i) Tcirg1 and (ii) Atp6v0a1, which encode the mammalian 116 kDa *a3* and *a1* subunits of the osteoclast vacuolar proton pump, respectively with mutations in *a3* alone accounting for over 50% of infantile malignant ARO (Frattini et al., 2000); (iii) Clcn7, a chloride voltage-gated channel indispensable for osteoclast function in mice and in humans (Kornak et al., 2001); (iv) Atp2b1, a P-type Ca^2+^-ATPase that regulates bone mass by fine-tuning osteoclast differentiation and survival (Kim et al., 2012); (v) Slc17a5, a lysosomal H^+^-driven sialic acid transporter whose mutations underpin Salla disease, a lysosomal storage disorder accompanied by bone malformations (Verheijen et al., 1999) and; lastly (vi) Slc37a2 (**Fig. 1C, red spot**), a little-studied member of the Slc37 family (isoforms a1-a4) (Cappello et al., 2018) that was enriched 2.35 Log_2_FC (*P=3.11E^-02^*) on SLs and whose human homologue harbors a genome-wide significant signal for eBMD (lead SNP rs7949048, *P*=3.1E^-12^) (**Fig. 1E, red box**), thus prompting our interest and further investigation.

### Slc37a2 is expressed in mature osteoclasts and localizes to a network of tubular SLs

To explore the relevance of Slc37a2 to osteoclasts and bone biology, we first assessed Slc37a2 expression in long bones using specific antibodies raised against mouse Slc37a2. As shown in **Fig. 1F**, Slc37a2 is highly expressed in mature osteoclasts, here lining endocortical bone, with immunofluorescent signal concentrated within F-actin rings as well as diffusely labelling the cell surface. In comparison, Slc37a2 expression is below the detection limit in neighbouring bone marrow, osteoblasts and osteocytes. Consistent with this expression pattern, Slc37a2 protein levels were robustly increased during RANKL-induced differentiation of BMMs, with peak expression observed at Day 5 when osteoclasts reach maturity (**Fig. 1G**). Thus, these findings imply that mature osteoclasts are the major, if not exclusive, source of Slc37a2 among bone-lineage cells.

Next, we examined the subcellular distribution of Slc37a2 in osteoclasts. First, we used immunoblotting to validate independently the enrichment of Slc37a2 during the biochemical isolation of SLs from mouse osteoclasts. In keeping with the proteomic data, Slc37a2 co-fractionates with major SL markers (as indicated here using LAMP2, V-ATPase subunit *a3* and Ctsk), with highest enrichment being observed in the SL fractions F1-F2 (**Fig. 2B**). To verify its localization to SLs in intact cells, we further immunostained primary cultures of mouse BMM-derived osteoclasts using the same Slc37a2-specific antibody along with the SL integral membrane protein LAMP2. As shown in **Fig. 2C,** Slc37a2 labelling resulted in a punctate staining pattern that overlapped with the pattern obtained with LAMP2 antibodies. Higher magnification revealed that Slc37a2 positive puncta colocalized with LAMP2, as indicated by the close overlap in fluorescent signal peaks in line scans (**Fig. 2D**).

**Fig. 2.**
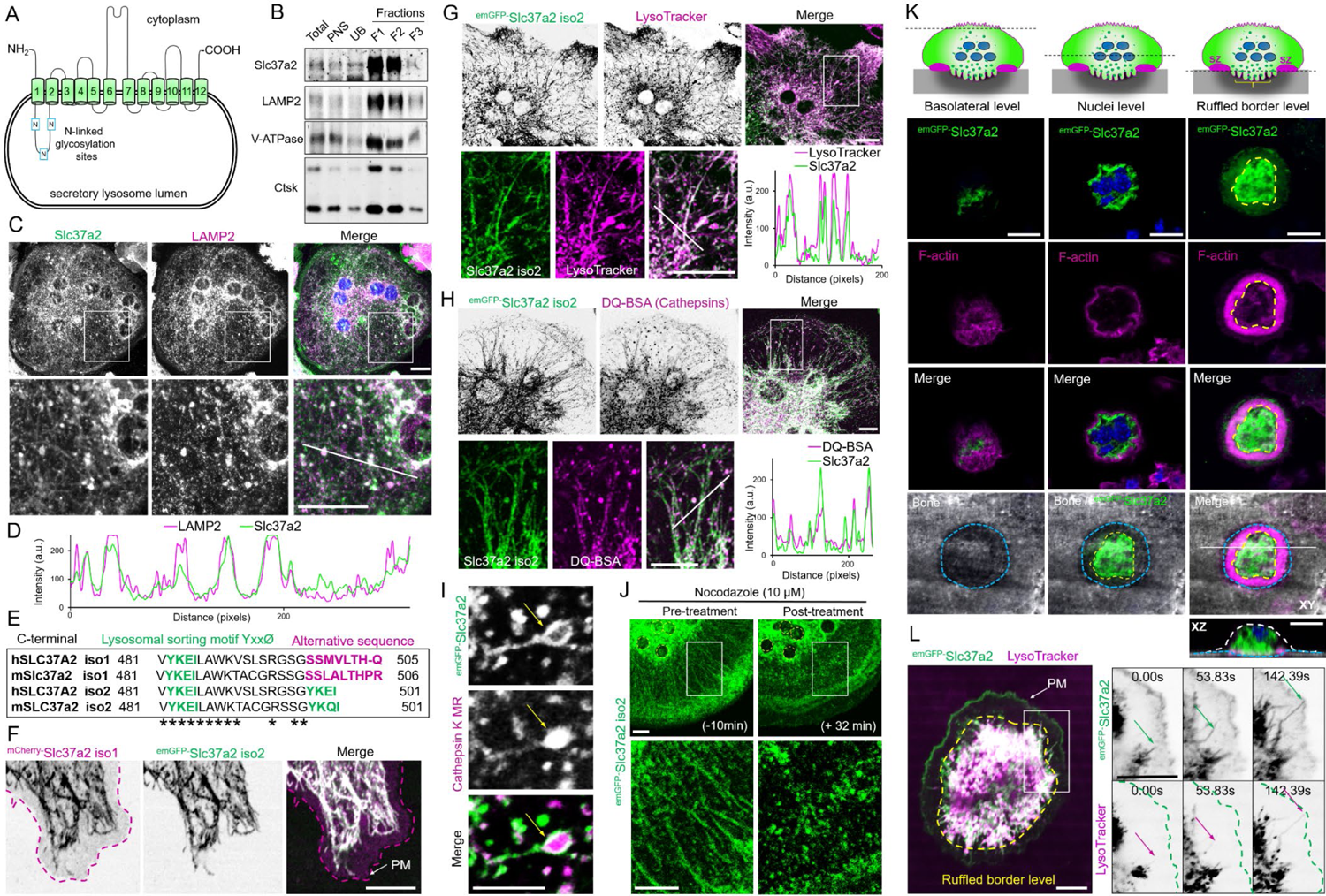
Slc37a2 localizes to a dynamic network of tubular SLs. (A) Schematic of Slc37a2 structure and topology on SL membranes, with transmembrane (green cylinders), luminal cytosolic domains (black lines) and N-linked glycosylation sites (blue boxes, N) indicated. (B) Slc37a2 co-enriches with SLs. 5 µg of proteins of indicated fractions collected during the isolation of enriched SLs was analyzed by SDS-PAGE and immunoblotting for Slc37a2 and SL markers LAMP2, V-ATPase and Ctsk. (**C-D**) Representative confocal image of a mouse BMM-derived osteoclast grown on glass and immunostained for Slc37a2 and LAMP2. White box corresponds with magnified regions. Bars, 10 µm (**C**). Line scan of white diagonal line in (**C**) indicating the spatial overlap in fluorescence signal intensities (**D**). (E) Amino acid alignment of the far C-terminus for human SLC37A2 and mouse Slc37a2 isoforms (1 & 2) with lysosomal sorting motifs and alternative sequences indicated. Asterisks indicate conserved amino acids. (F) Confocal image of a mouse osteoclast cultured on glass co-microinjected with ^mCherry-^ Slc37a2 isoform 1 and ^emGFP-^Slc37a2 isoform 2. Purple dashed line outlines the plasma membrane (PM). Bar, 10 µm. (**G-H**) Representative confocal image of a live mouse osteoclast on glass microinjected with ^emGFP-^Slc37a2 isoform 2 and pulsed with endolysosomal probes LysoTracker Red (**G**) or DQ-BSA (**H**). Line scans of the individual fluorescent channels correspond to the white diagonal line in the magnified views. Bars, 10 µm. (I) High-resolution live confocal image of a representative tubular SL bearing ^emGFP-^Slc37a2 isoform 2 and housing Cathepsin K Magic Red (MR) within its lumen (yellow arrow). Bar, 2 µm. (J) Time-lapse confocal image of an osteoclast expressing ^emGFP-^Slc37a2 isoform 2 before (−10 min) and after (+ 32 min) treatment with the microtubule disrupting agent nocodazole (10 µM). (K) Multilevel confocal images of a mouse osteoclast cultured on bone expressing ^emGFP-^ Slc37a2 isoform 2 and stained with rhodamine-phalloidin to indicate the F-actin ring and underlying ruffled border. Representative XY serial confocal sections depict ^emGFP-^Slc37a2 distribution at the basolateral, nuclei (blue) and the ruffled border level. XZ denotes side view corresponding to the white horizontal line in the merged panel (lower left). Dashed lines indicate ruffled border region within sealing zones (yellow) and resorptive pit (blue). Bars, 10 µm. (L) Bottom-up time-lapse confocal image of an osteoclast cultured on bone expressing ^emGFP-^ Slc37a2 isoform 2 and labeled with LysoTracker Red. The magnified region represents a time-lapse series illustrating SL tubulation and fusion with the bone-apposed plasma membrane, with time points indicated. Dashed line circumscribes the sealing zone (yellow) and bone-orientated membrane. Arrows track the dynamics of Slc37a2 (green) and LysoTracker (purple), respectively along the same extending tubular SL. Bars, 10 µm.

Among the Slc37 family, Slc37a2 is unique, existing as an N-terminal glycosylated 12-transmembrane spanning protein (**Fig. 2A**) encoded by two naturally occurring splice variants (isoforms 1 & 2) that differ by five amino acids found within their extreme C-terminus. Slc37a2 isoform 2 harbors a canonical lysosomal sorting signal (YxxØ) whereas isoform 1 possesses an extended alternative sequence SSxxxLTH-x that is conserved both in humans and in mice (**Fig. 2E**). Therefore, to differentiate the subcellular distribution of the two Slc37a2 isoforms in osteoclasts, the N-terminus of each mouse Slc37a2 variant was tagged with either emGFP (isoform 2) or mCherry (isoform 1) and cloned into a CMV-driven mammalian vector. Because osteoclasts are refractory to conventional transfection methods, tagged-Slc37a2 variants were co-expressed by direct nuclear microinjection into mouse BMM-derived osteoclasts. Remarkably, when microinjected into live osteoclasts and imaged by time-lapse confocal microscopy on glass, Slc37a2 isoforms co-occupied an expansive network of highly dynamic tubulo-vesicular compartments that radiated throughout the osteoclast cytoplasm (**Fig. 2F-J, Video S1**). As shown in **Fig. 2F**, whereas the subcellular distribution of ^emGFP-^Slc37a2 isoform 2 tightly overlapped with ^mCherry-^Slc37a2 isoform 1 on tubular organelles, Slc37a2 isoform 1 showed additional preference for the plasma membrane (PM, purple outline), likely reflecting its alternative C-terminal targeting signal. Interestingly, the lumens of these tubular organelles were acidic and housed cathepsins/K (**Video S1)**, as confirmed by the accumulation of the acidophilic probe LysoTracker Red (**Fig. 2G**) and cathepsin/k fluorescent substrates DQ-BSA (pan-cathepsins) (**Fig. 2H**) and Magic Red (cathepsin K MR) (**Fig. 2I**), indicating that they were of endolysosomal origin. This was corroborated by monitoring co-trafficking of ^emGFP-^ Slc37a2^+^ organelles with ^mRFP-^LAMP1 as well as LRO marker proteins ^mCherry-^VAMP7 and ^mCherry-^Rab38 in living osteoclasts (**Extended Data Fig. 1A-B, Video S1**). By comparison, while Slc37a2^+^ tubules closely intersected with other organelles including early endosomes (^mCherry-^ Rab5) and mitochondria (Mitotracker Red), they were morphologically and dynamically distinct (**Extended Data Fig. 1A-B and Video S2).**

The Slc37a2^+^ tubular lysosomal network was highly sensitive to chemical and cold fixation (i.e. 4% PFA) and rapidly collapsed upon treatment with the destabilizing reagent nocodazole (**Fig. 2J**), indicating that it is intimately linked to microtubules (Swanson et al., 1987). In addition, we frequently observed fusion of Slc37a2^+^ tubules with the osteoclast plasma membrane (**Video S3**). In keeping with the view that the ruffled border is formed by the insertion of SLs to the bone-apposed plasmalemma (Stenbeck, 2002), when cultured on devitalized bone surfaces, ^emGFP-^Slc37a2^+^ tubules concentrated within F-actin rings demarcating ruffled borders (**Fig. 2K**) and fused with the ventral plasma membrane when monitored under live settings following lentiviral-mediated delivery of ^emGFP-^Slc37a2 (**Fig. 2L arrows, Video S4**). Importantly, these tubular compartments were not an artefact of ^emGFP-^ Slc37a2 expression, as the same network of acidified, cathepsin-containing tubular organelles could be visualized in naive osteoclasts using vital probes LysoTracker and/or DQ-BSA by optical (**Extended Data Fig. 2A-B, Video S5**) and by electron (**Extended Data Fig. 2C**) microscopy. Collectively, these data indicate that Slc37a2 occupies a network of acidified cathepsin-containing tubular LROs that fuse with the bone-apposed plasmalemma, thus fulfilling the usual criteria of osteoclast SLs. Herein, we refer to Slc37a2^+^ compartments as ‘tubular SLs’.

### *Slc37a2* deficiency leads to high bone mass

To investigate the physiologic importance of the Slc37a2^+^ tubular SL network in osteoclasts and bone metabolism, we next generated Slc37a2 knockout (KO) mice using targeted embryonic stem cells obtained from the ‘Knockout Mouse Project’ consortium. Genotyping and qPCR analysis confirmed correct insertion of the tma2 targeting cassette and that the approach produced an effective knockdown of the *Slc37a2* mRNA (>95% reduction in *Slc37a2*) in bones from homozygous mice compared to wild-type (WT) littermates (**Extended Data Fig. 3A-C**). *Slc37a2^tm2a(KOMP)wtsi^* homozygous mice, referred herein as “*Slc37a2* knockout” (*Slc37a2*^KO^) mice, are born at Mendelian frequency and are indistinguishable from WT littermates at birth (data not shown). No overt differences in the size or weight of matched littermates were observed up to 24-weeks-of-age (**Extended Data Fig. 3D**). Similarly, no obvious abnormality in skeletal patterning was observed in whole mount alcian blue/alizarin red staining of skeletal preparations of 5-day-old *Slc37a2*^KO^ mice (**Extended Data Fig. 3E**) or by whole body mammography of 12-week-old female mice (**Extended Data Fig. 3F**). On the other hand, radiographs and microcomputed tomography (µCT) analysis of the distal femurs (**Fig. 3A-D**) of 12-week-old WT and *Slc37a2*^KO^ mice revealed that the long bones of null mice were radiodense and exhibited a dramatic increase in trabecular bone volume; reaching an impressive ∼300-1000% increase in BV/TV in male and female mice, respectively (**Fig. 3E**). Consistent with this finding, femurs of *Slc37a2*^KO^ mice had increased trabeculae and a decrease in trabecular separation (**Fig. 3F-G**), but only females showed a significant increase in trabecular thickness at 12-weeks (**Fig. 3H**). Cortical bone thickness was also significantly increased in both male and female mice at 12-weeks (**Fig. 3D & I**) which imparted greater bone stiffness and whole bone strength as confirmed by three-point-bending tests (**Fig. 3J-K**). Elevated bone mass was also evident in heterozygous (*Slc37a2*^HET^) mice, indicative of a gene dose effect (**Extended Data Fig. 3G,J-N**). This profound increase in bone mass extended to vertebrae within the axial skeleton (**Fig. 3L-N**) as well as to the bones of the skull, albeit to a lesser extent (**Extended Data Fig. 3H**). While femur length was unaltered (**Extended Data Fig. 3I**), trabecular bone mass accumulated and was retained with age (**Fig. 3O-Q**).

**Fig. 3.**
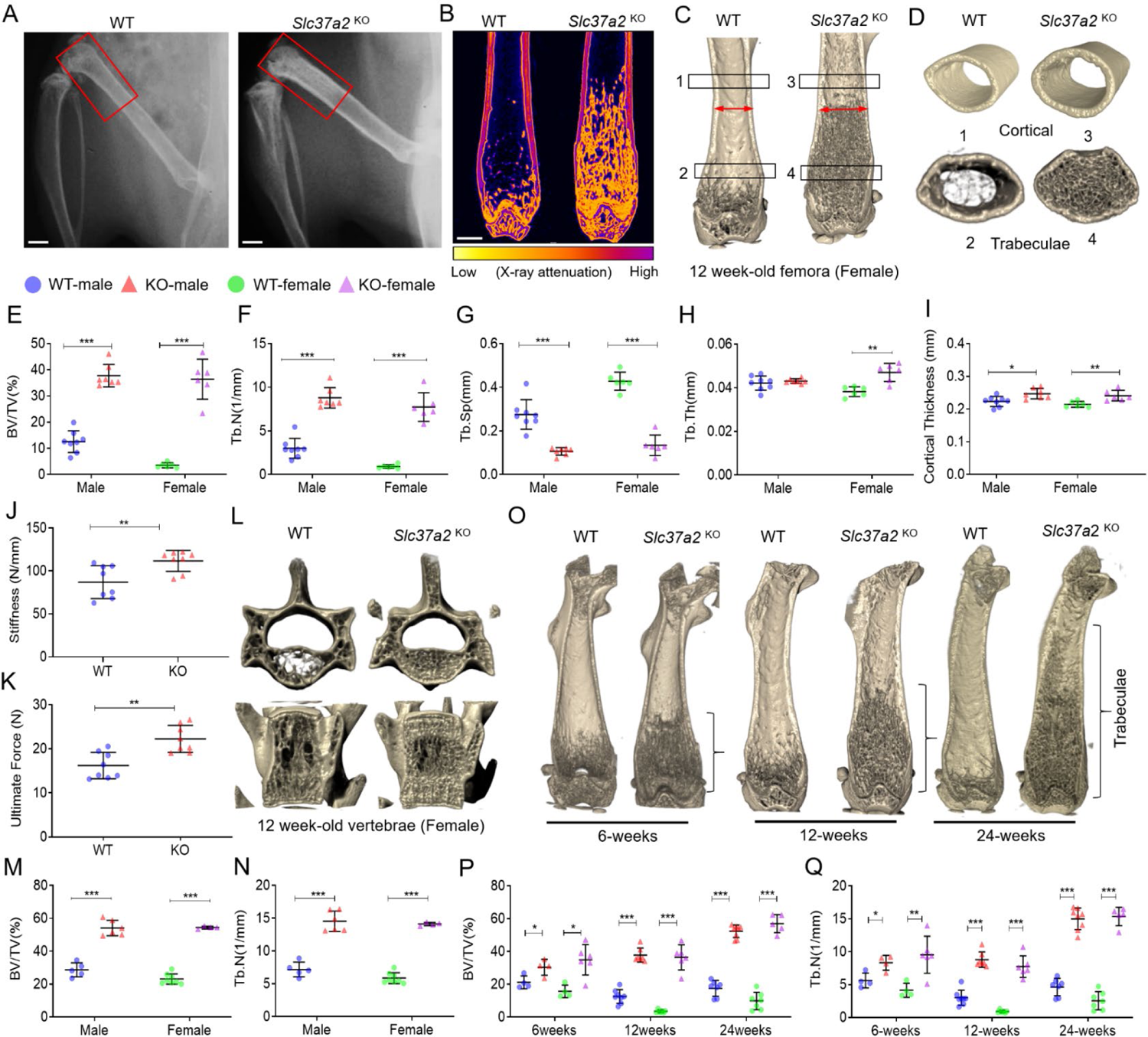
*Slc37a2*^KO^ exhibit high bone mass and increased bone strength. (**A**) Radiography of the hindlimbs of 12-wk-old female WT and *Slc37a2*^KO^ mice. Bar, 1 mm. (**B-D**) Representative sagittal µCT section false colored to indicate X-ray attenuation (**B**) and µCT 3D reconstructed images of sagittal (**C**) and transverse (**D**) sections of distal femurs of 12-wk-old female WT and *Slc37a2*^KO^ mice. (**E-I**) µCT analysis of femurs of 12-wk-old male and female WT and *Slc37a2*^KO^ mice (*n =* 6-8). (**J-K**) Biomechanical three-point-bending test of 12-wk-old male WT and *Slc37a2*^KO^ femurs. Stiffness (**J**) and Ultimate Force (**N**) are presented. (*n* = 8). (**L-N**) Representative µCT images of vertebrae (**L**) and µCT analysis of vertebrae (**M-N**) of 12-week-old male and female WT and *Slc37a2*^KO^ mice. (*n =* 5–6). (**O-Q**) Representative µCT images of female femurs (**O**) and µCT analysis of femurs (**P-Q**) of 6-, 12- and 24-week-old male and female WT and *Slc37a2*^KO^ mice. (*n* = 4–7). All data are presented as means ±SD. *, *P* <0.05, ** *P*<0.01, *** *P* <0.001, by two-tailed unpaired Student’s t-test.

Histologically, *Slc37a2*^KO^ femurs contained cartilaginous bars deep within trabeculae of the metaphysis (**Fig. 4A, inset**), suggestive of impaired calcified cartilage degradation during endochondral bone growth. Modeling of *Slc37a2*^KO^ mice bones was also abnormal as evidenced by the broadening of the distal diaphysis and metaphysis (**Fig. 4B, dual-arrows**). Moreover, femurs of *Slc37a2*^KO^ mice exhibited a conspicuous fibro-cartilaginous lesion that occurred at the level of the periosteum and extended along the proximal metaphyseal-diaphyseal junction (**Fig. 4C-H**). This lesion stained intensely for TRAP activity, a marker of osteoclasts (**Fig. 4E**) and was interspersed with small TRAP^+^α_v_β^+^ cells that lined the periosteal bone surface as well as numerous bone/cartilage fragments (**Fig. 4E-F, arrows**) that appeared partially demineralized under transmission electron microscopy (**Fig. 4G, arrows**). In addition, this periosteal lesion was densely populated with Runx2^+^ mesenchymal cells that stained strongly for MMP13, a matrix metalloprotease known to participate in the digestion of cartilage and collagen matrix (**Fig. 4H**). Outside of the skeleton, macroscopic and histopathological evaluation of major organs and tissues revealed no obvious abnormalities in *Slc37a2*^KO^ mice at 12-weeks (**Extended Data Fig. 3 O-P**). Taken together, these data suggest that the major phenotype of mice with global *Slc37a2* insufficiency is high bone mass.

**Fig. 4.**
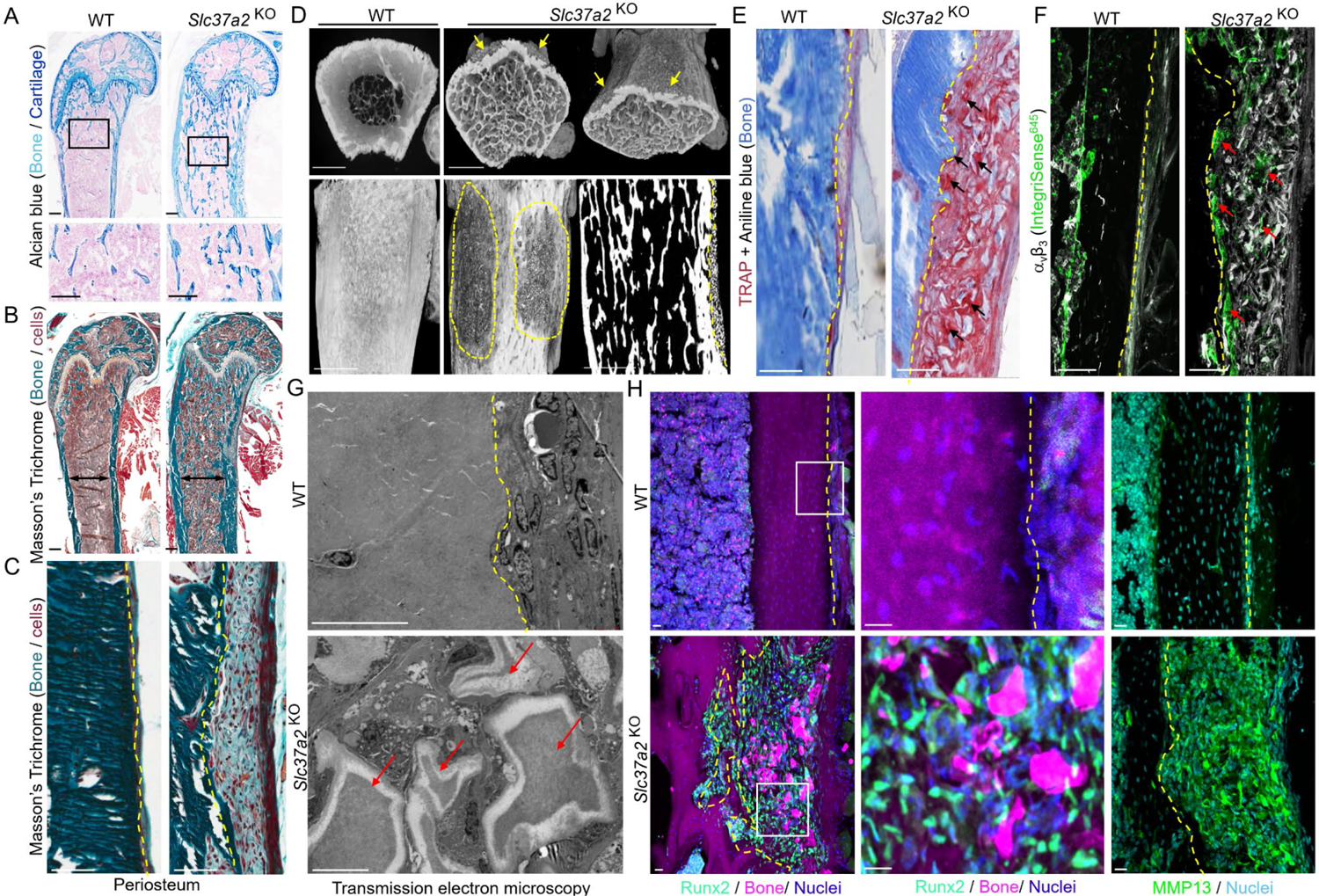
*Slc37a2*^KO^ mice exhibit high bone mass and a periosteal bone lesion. (A) Representative histological images of Alcian blue staining of distal femur of 12-week-old female WT an*d Slc37a2*^KO^ mice, Bars, 0.1 mm. (**B, C**) Mason’s Trichome staining of femurs from 12-week-old female WT and *Slc37a2*^KO^ mice (B) and representative images of the periosteal surface along the metaphyseal-diaphyseal junction (**C**). Black double-arrows indicate maximal femoral diameter and dashed line indicates the periosteal bone surface. Bars, 1 mm. (**D**) High-resolution µCT images of 12-week-old female WT and *Slc37a2*^KO^ femurs indicating the periosteal lesions (arrows and dashed outlines). Bars, 1 mm. (**E-F**) Histological evaluation of TRAP^+^ (**E**) and αvβ3^+^ (**F**) cells along the periosteal surface of 12-week-old femurs from female WT and *Slc37a2*^KO^ mice. Dashed line denotes periosteal bone surface, arrows indicate TRAP^+^ or αvβ3^+^ cells. (G) Transmission electron microscopic evaluation of the periosteal layer of femurs from WT and *Slc37a2*^KO^. Dashed line indicates periosteal bone surface. Red arrows depict demineralized bone/cartilaginous fragments. Bars, 10 µm. (H) Representative images of cryosections depicting the periosteal surface along the femurs of 12-week-old female WT and *Slc37a2*^KO^ mice immunostained with Runx2 or MMP13.

### High bone mass in *Slc37a2*^KO^ mice is driven by impaired bone resorption and compensatory increases in remodeling-based bone formation

To determine the cellular mechanism(s) by which *Slc37a2*-deficiency increased trabecular bone volume, histomorphometric and fluorescence immunohistochemical analysis of the trabecular surfaces of the femur were performed. First, osteoclast parameters were assessed and revealed a dramatic increase in both the total number of TRAP^+^ osteoclasts (∼246%, *P*<0.0001) and number of osteoclasts lining trabecular bone surfaces (∼182%, *P*<0.0001) of *Slc37a2*^KO^ mice by histomorphometry (**Fig. 5A-C**). This increase in osteoclast numbers was corroborated by elevated systemic levels of TRAP5b (**Fig. 5D**) as well as increased expression of key osteoclast marker genes Acp5 and Ctsk in femurs (**Fig. 5P**). Reflecting this, RANKL mRNA was robustly amplified in *Slc37a2*^KO^ femurs, whereas OPG and RANK mRNA levels were unchanged, leading to a significant increase in the mRNA ratio of RANKL-to-OPG (**Fig. 5E**). *In situ* assessment of trabecular surfaces revealed that whereas *Slc37a2*^KO^ osteoclasts were capable of forming F-actin rings, they were larger than WTs (**Fig. 5F-H**). Despite this, serum C-terminal telopeptides of type I collagen (CTX-I), a by-product of collagen type-1a that is released from bone into blood by osteoclast resorption, was not significantly altered in *Slc37a2*^KO^ mice (**Fig. 5I**). However, the ratio between CTX-I and TRAP5b, which is used as an index for resorption per osteoclast (Henriksen et al., 2007), revealed a highly significant decrease in resorption per osteoclast (*P*=0.0006) (**Fig. 5J**). Intriguingly, resorptive lacunae underlying αvβ3^+^ *Slc37a2*^KO^ osteoclasts were frequently occupied by flattened bone-lining MMP13^+^ cells (**Fig. 5K, arrows**), in keeping with the view that bone lining ‘reversal’ cells play an accessory role in removing residual demineralized bone matrix left by osteoclasts (Everts et al., 2002, Andersen et al., 2004, Pirapaharan et al., 2019). Along with increased expression of MMP13, the mRNA levels of MMP9 and MMP14 were also significantly elevated in *Slc37a2*^KO^ femurs (**Fig. 5P**), inferring that the impaired bone resorption in *Slc37a2*^KO^ mice is partially compensated by increased expression of proteases.

**Fig. 5.**
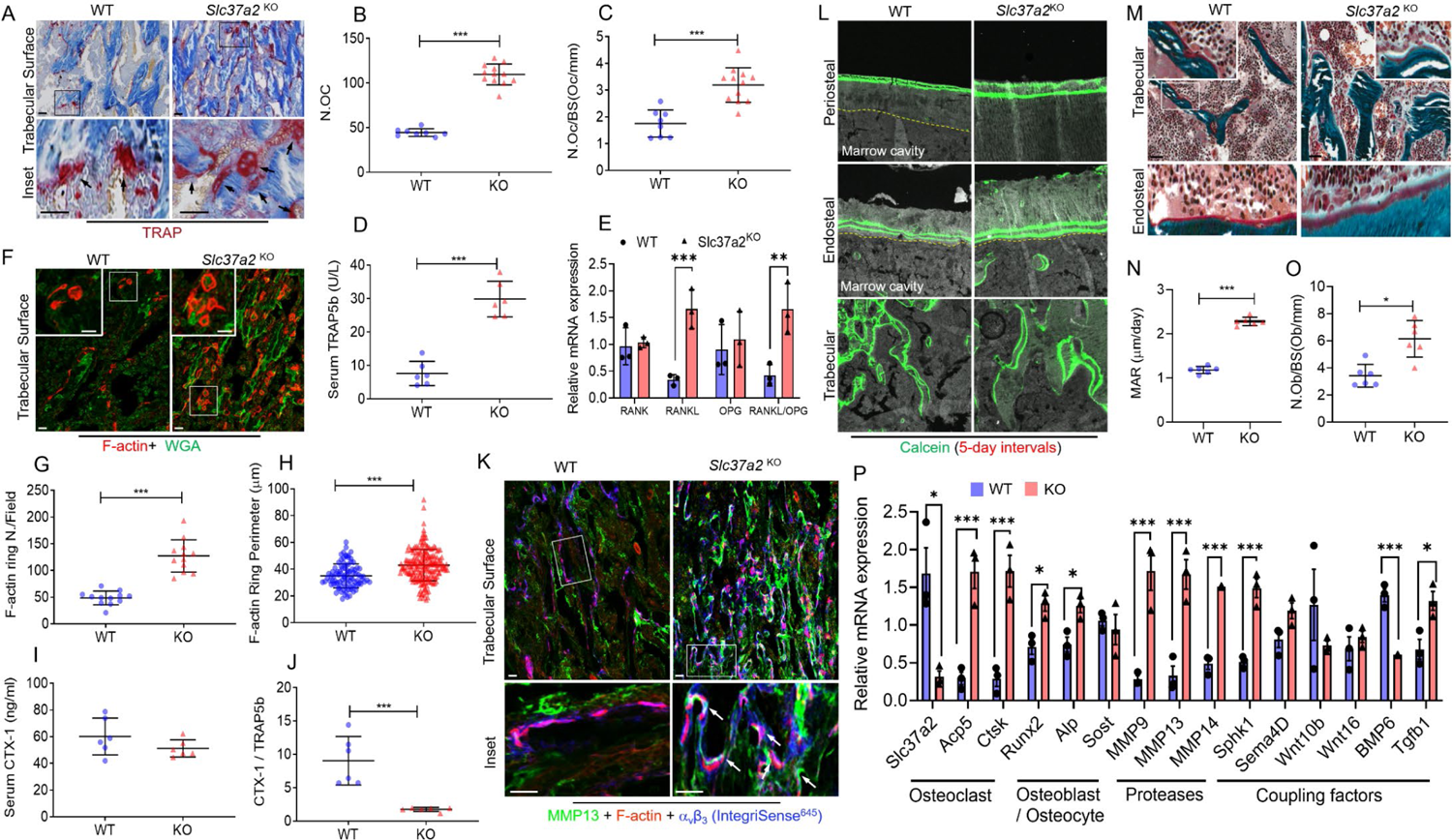
*Slc37a2* deletion impairs bone resorption leading to compensatory increases in osteoclastogenesis, protease expression and osteoblast-mediated bone formation. (**A**) Representative images of the primary spongiosa of proximal WT and *Slc37a2*^KO^ male femora stained for TRAP and (**B-C**) histomorphometric analysis of TRAP^+^ osteoclasts (*n* = 8-12). Bar, 50 μm. (**D-E**) Serum TRAP5b levels (**D**, *n* = 6) and RANK/RANKL/OPG mRNA expression in femurs (**E**, *n* = 3) of 12-week-old male WT and *Slc37a2*^KO^ mice. (**F-H**) Confocal images of representative femur cryosections stained for F-actin and Wheat-germ agglutinin, (WGA) from 12-week-old male mice (**F**) and quantitation of F-actin numbers (**G**, *n* = 12 fields/group) and diameters (**H**, *n* = 92-192 cells/group). Bars, 50 μm. (**I-J**) CTX-1 levels (**I**) and ratio of CTX-1/TRAP5b levels (**J**) in serum of 12-week-old male WT and *Slc37*^a2^ mice (*n* = 6). (**K**) Representative confocal image of the primary spongiosa of WT and *Slc37a2*^KO^ male femora immunostained for MMP13, F-actin and αvβ3 (IntegriSense^645^). Arrows indicate flat bone-lining MMP13^+^ cells occupying resorptive lacunae in magnified images. Bars, 50 μm. (**L-O**) Representative images of 12-week-old male femurs from WT and *Slc37a2*^KO^ mice depicting fluorescent calcein-double labelling (**L**), Mason’s Trichrome staining (**M**) and histomorphometric analyses of bone mineral apposition rate (**N**) and osteoblast numbers/ bone surface (**O**) (*n* = 6). (**P**) Relative mRNA expression in femora from 12-week-old WT and *Slc37a2*^KO^ mice (*n* = 3). All data are presented as means ± SD. * *P* < 0.05, ** *P* < 0.01, \*\*\**P* < 0.001 by two-tailed unpaired Student’s t tests.

Because osteoclast-mediated bone resorption is intimately coupled to bone formation by osteoblasts (Sims and Martin, 2020), we also examined bone formation parameters by dynamic bone histomorphometry of double calcein-labeled frozen sections (**Fig. 5L**). As shown in **Fig. 5N**, femora isolated from *Slc37a2*^KO^ mice exhibited significantly increased mineral apposition rates (MARs) compared with WT. Consistently, the number of osteoblasts occupying bone surfaces (**Fig. 5M,O**), as well as the expression of osteoblast marker genes Runx2 and ALP (**Fig. 5P**), were increased in distal femur metaphysis of 12-week old *Slc37a2*^KO^ mice. The capacity of *Slc37a2*^KO^ osteoblasts to differentiate and form bone mineralizing nodules *in vitro*, however, was indistinguishable from WT osteoblasts (**Extended Fig. 4),** implying that the observed increase in bone formation *in vivo* was indirect. Corroborating this, analyses of the mRNA expression of key osteoclast-derived ‘clastokines’ (Sphk1/S1P, Sema4D, Wnt10b, Wnt16 and BMP6) and the matrix-derived osteoblast ‘coupling factor’ (TGF-β) (Sims and Martin, 2020) in femurs revealed that loss of Slc37a2 in mice corresponded with a significant increase in Sphk1 and TGF-β1 mRNA expression, but not Sem4A (*P*=0.06) Wnt16, Wnt10b or BMP6 (**Fig. 5P**). Thus, high bone mass in *Slc37a2*^KO^ mice is driven, primarily, by impaired bone resorption by osteoclasts together with imbalanced remodeling-based bone formation by osteoblasts.

### *Slc37a2* deletion impairs the function but not formation of osteoclasts

We next assessed whether Slc37a2 regulates cell-autonomous effects in osteoclasts. To this end, BMMs isolated from WT and *Slc37a2*^KO^ littermates were exposed to M-CSF and RANKL for 5 days to induce osteoclast differentiation. As shown in **Fig. 6A-B**, the number of TRAP^+^ osteoclasts formed *in vitro* is indistinguishable between WT and *Slc37a2*^KO^ mice. Consistently, no obvious morphological distinctions were observed in either the spreading or capacity of *Slc37a2*^KO^ osteoclasts to form F-actin rich podosome-belts (**Fig. 6C**). Temporal expression of protein markers for osteoclast differentiation, including NFATc1, c-Src, V-ATPase or cathepsins K and B, were also comparable between WT and *Slc37a2*^KO^ cells (**Fig. 6D**). Further, absence of *Slc37a2* did not alter key RANKL-induced signaling events, including phosphorylation of ERK1/2, Akt and IκBα (**Fig. 6E**). Thus, Slc37a2 does not regulate osteoclast differentiation.

**Fig. 6.**
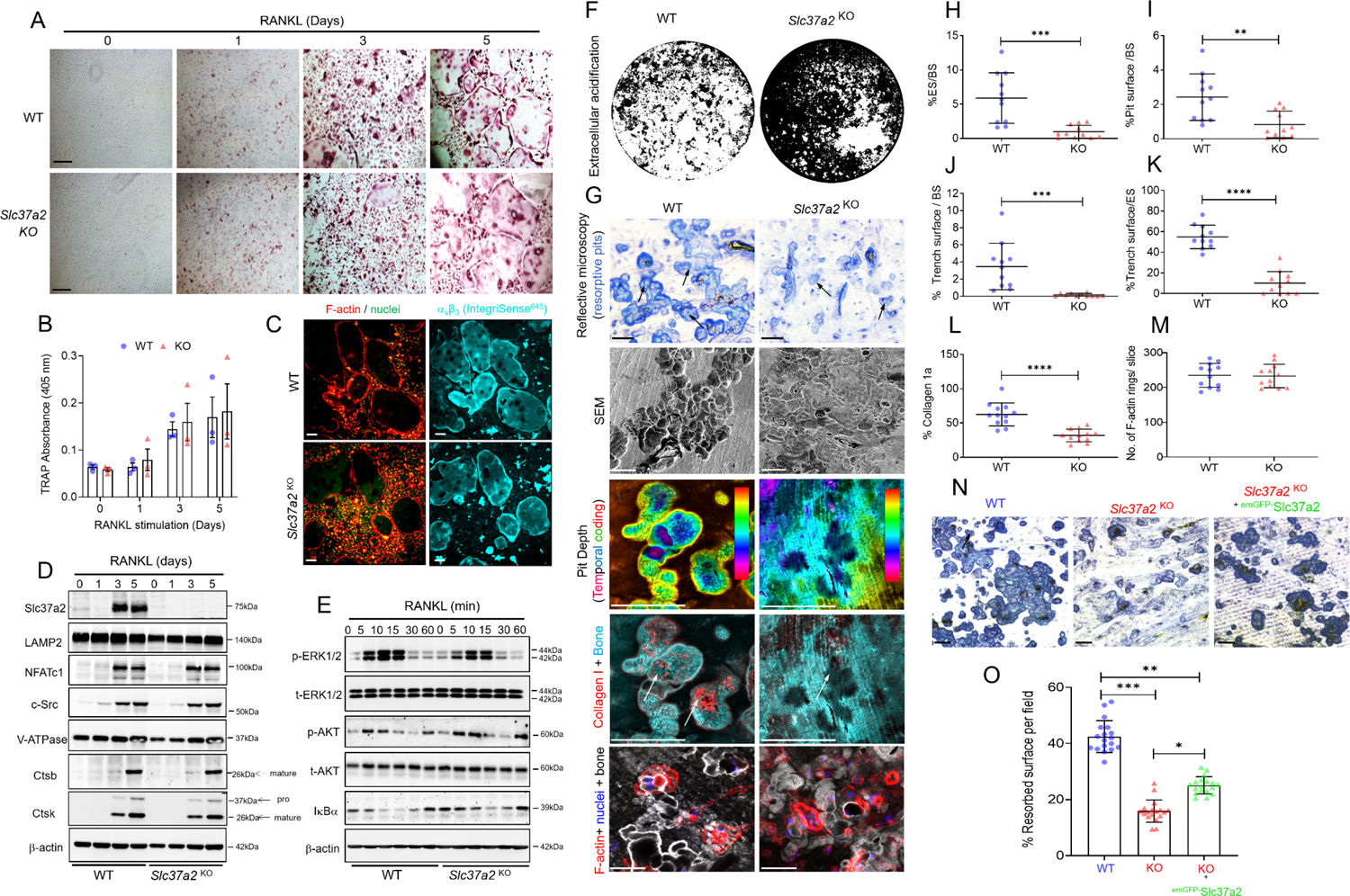
*Slc37a2* deletion impairs bone resorption but does not alter osteoclast differentiation, spreading or RANKL-signaling cascades. (**A-B**) WT and *Slc37a2^KO^* BMMs were cultured with M-CSF (25 ng/ml) and RANKL (10 ng/ml) and multinucleated cells were TRAP stained (**A**), and TRAP activity was then measured by colorimetric analysis (405 nm) (**B**) (*n* = 3). Bar, 50 µm. (C) Representative confocal images of WT and *Slc37a2^KO^* osteoclasts cultured on glass and immunostained for F-actin, αvβ3 (IntegriSense^645^) and nuclei. Bar, 50 µm. (D) Expression of osteoclast and SL marker proteins during RANKL-induced differentiation of WT and *Slc37a2^KO^* BMMs by immunoblotting using indicated antibodies. (E) RANKL-signaling in WT and *Slc37a2^KO^* BMMs. Immunoblotting was used to monitor the levels and phosphorylation states of ERK1/2, AKT and IκBα in response to RANKL. (F) Representative image of Von Kossa staining of WT and *Slc37a2^KO^* osteoclasts cultured on mineralized substrates. White clearings denote resorbed mineral. (**G-M**) Bone resorptive activity in *Slc37a2^KO^* osteoclasts is impaired. Shown are representative images of *in vitro* bone resorption assays of WT and *Slc37a2^KO^* osteoclasts visualized by reflective microscopy, SEM, and confocal microscopy with quantification of percentage of eroded surface (**H**), pit area (**I**), trench surface per bone slice (**J**), trench surface/eroded surface (**K**), collagen1a staining (**L**) and number of F-actin rings per bone slice (**M**) Black and white arrows indicate bone resorption pits in toluidine blue and collagen 1 stained bone slices, respectively. (*n* = 11-12 bone slices pooled from 3 independent experiments per genotype). Bars, 50 µm. (**N-O**) Rescue of *Slc37a2^KO^* osteoclasts activity by lentiviral mediated transduction of ^emGFP-^ Slc37a2. Depicted are representative images of toluidine blue stained resorptive pits of WT, *Slc37a2^KO^* and ^emGFP-^Slc37a2 transduced *Slc37a2^KO^* osteoclasts with quantification of the percentage resorbed surface per field quantified (*n* = 54 fields from 3 experimental replicates). All data are presented as means ± SD. * *P* < 0.05, ** *P* < 0.01, *** *P* <0.001 by two-tailed unpaired Student’s t test (**H-M**) or nested one-way ANOVA (**O**). Bar, 50 µm.

In comparison, when cultured on synthetic bone mineral (**Fig. 6F**) or bovine bone slices (**Fig. 6G**), *Slc37a2*^KO^ osteoclasts exhibited dramatically diminished capacity to acidify their extracellular environment and digest mineralized matrix. Unlike the well-demarcated resorption lacunae excavated by WTs, resorption pits generated by *Slc37a2^KO^* osteoclasts were morphologically irregular, formed fewer ‘trench-like’ resorptive trails and were visibly shallow as revealed by reflective and scanning electron microscopy (**Fig. 6G-L**). Confirming this, projection of confocal z-stacks of representative resorptive pits revealed shallower temporal depth and reduced collagen Type 1a exposure in pits formed by *Slc37a2*^KO^ osteoclasts compared to WTs (**Fig. 6L**), reflecting their reduced extracellular acidification function. Intriguingly, this resorption deficit was independent of their capacity to form F-actin rings/sealing zones (**Fig. 6M**). Moreover, the pit phenotype could be rescued by transduction of recombinant lentivirus containing ^emGFP-^Slc37a2 into osteoclasts derived from *Slc37a2*^KO^ BMMs (**Fig. 6N-O**), confirming that the impairment was Slc37a2-dependent. Together, these data indicate that Slc37a2 regulates the function, but not the differentiation, of osteoclasts in a cell-autonomous way.

### Slc37a2 deficiency impairs the export of monosaccharide sugars from SLs required for delivery to the ruffled border

To investigate the mechanism(s) by which Slc37a2 regulates osteoclast activity, comparative multi-omics analyses were performed on osteoclasts from WT and *Slc37a2*^KO^ mice (**Extended Data Fig. 5**). Except for the notable absence of Slc37a2, no consistent changes were detected both at the mRNA and protein levels between *Slc37a2*^KO^ and WT osteoclasts as determined by either RNAseq or label-free quantitative proteomics using a minimum of >1.5 Log_2_FC and an adjusted *P* value of <0.05 (**Extended Data Fig.5**). Similarly, the proteomes of SLs isolated from *Slc37a2*^KO^ and WT osteoclasts were unremarkable (**Extended Data Fig. 5**). This suggests that Slc37a2 deficiency does not alter the global transcriptome or proteome of osteoclasts. On the other hand, quantitative metabolomic profiling of osteoclasts derived from WT and *Slc37a2*^KO^ mice revealed differences in the levels of several key metabolites, most notably monosaccharide sugars including D-(+)-glucose 1 and D-(-)-fructose1/2, which were increased in *Slc37a2*^KO^ osteoclasts compared to WT (**Fig. 7A**). The level of glucose-6-phosphate (G6P), the presumed substrate of Slc37a2 (Cappello et al., 2018), was however unaltered in *Slc37a2*-deficient osteoclasts.

**Fig. 7.**
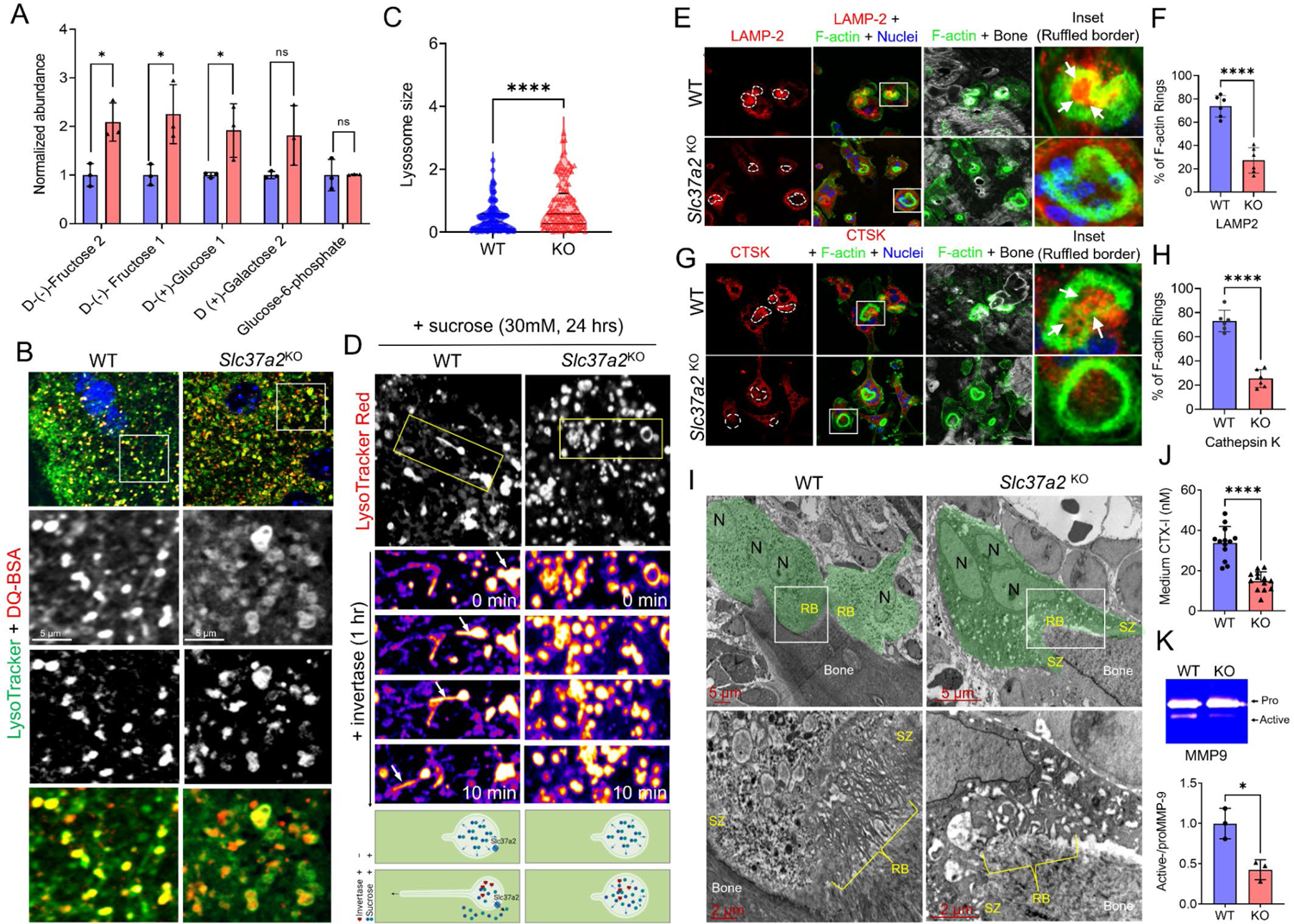
Export of monosaccharides and delivery of SLs to the ruffled border is impaired. *Slc37a2*^KO^ osteoclasts. (**A**) Quantitative metabolomics profiling of monosaccharide sugars in BMM-derived osteoclasts from WT and *Slc37a2^KO^* mice (*n* = 3). (**B-C**) Representative confocal images of WT and *Slc37a2^KO^* osteoclasts stained with LysoTracker Green and DQ-BSA (**B**) and quantitation of SL size (**C**). (*n* = 109 SLs pooled from 10 cells per group). (**D**) Time-lapse confocal image series of WT and *Slc37a2^KO^* osteoclasts cultured in high sucrose conditions (30 mM, 24 h) to induce sucrosome formation and imaged for 10 min post-treatment with invertase (0.5 mg/ml, 1h). SLs are were pulsed with LysoTracker Red prior to imaging. Arrows track a SL resolution and tubulation event as summarized in the underlying cartoon. (**E-H**) Confocal images of WT and *Slc37a2^KO^* osteoclasts cultured on bone and immunostained for F-actin in combination with either LAMP-2 (**E**) or cathepsin k (CTSK) (**F**) and quantitation (**G-H**). Arrows indicate delivery of LAMP2 and CTSK within F-actin rings. (*n* = 6). (**K**) Representative TEM micrographs of WT and *Slc37a2^KO^* osteoclasts (green) lining trabecular bone surfaces within the primary spongiosa of 5-day-old male littermates. Magnified picture illustrate ruffled borders (RB). N=Nuclei, SZ=sealing zone. **(J-K**) CTX-1 and active MMP9 levels in media from cultures of bone-resorbing WT and *Slc37a2*^KO^ osteoclasts as monitored by ELISA (**J**) and gel zymography (**K**). *n* = 13 (**J**) and *n* = 3 (**K**). Data are presented as means ± SD * *P* <0.05 and **** *P* < 0.0001 by two-tailed unpaired Student’s t test.

It has long been known that glucose and fructose are transported out of lysosomes by an as yet unidentified monosaccharide transporter(s) (Mancini et al., 1990, Lizak et al., 2019). Given that Slc37a2 resides on osteoclast SLs and that its absence coincided with increased intracellular levels of glucose and fructose, we considered the possibility that Slc37a2 regulates the export of monosaccharide sugars out of SLs. According to this scenario, SLs in *Slc37a2*^KO^ osteoclasts should be larger than those in controls due to osmotically imposed swelling upon substrate accumulation. Consistent with this premise, the size of SLs (co-labeled with LysoTracker green and DQ-BSA) in *Slc37a2*^KO^ osteoclasts were dramatically increased, and associated network less tubular, when compared with their WT counterparts (**Fig. 7B-C**). Morphologically, *Slc37a2*^KO^ SLs appeared ‘swollen’ and resembled ‘sucrosomes’, (i.e. osmotically swollen endolysosomes that form upon the uptake and accumulation of undigestible sucrose) (Cohn and Ehrenreich, 1969). Addition of invertase, an enzyme that breaks down sucrose into its monosaccharide glucose and fructose constituents, is known to induce the resolution (shrinkage) of sucrosomes, back to lysosomes, a process that coincides with extensive tubulation of lysosomal membranes (Swanson et al., 1986, Bright et al., 2016). Consistently, addition of invertase (0.5 mg/ml, 1hr, *S. cerevisiae*) to osteoclast cultures superinfused with sucrose-containing solution (30 mM, 24 hrs) prompted rapid shrinkage and tubulation of WT LysoTracker Red^+^ sucrosomes (**Fig. 7D**), but failed to resolve sucrosomes in *Slc37a2*^KO^ osteoclasts when monitored over the same period by time-lapse confocal microscopy (**Video S6**). Thus, these data imply that Slc37a2 facilitates the export of glucose and fructose out of osteoclast SLs, a necessary prerequisite for the maintenance of SL size and the initiation of membrane tubulation.

In addition to initiating endomembrane remodeling events (Saric and Freeman, 2021), the efflux of luminal solutes is an important requirement for lysosomal transport (Bandyopadhyay et al., 2014). Therefore, we next asked whether the morphological disturbances observed in *Slc37a2*^KO^ SLs altered their capacity to translocate to the osteoclast ruffled border. To this end, we employed immunofluorescent confocal microscopy to monitor the delivery of key SL marker proteins LAMP2 and Ctsk to the ruffled border in bone-resorbing osteoclasts. As shown in **Fig. 7E-H**, whereas WT osteoclasts exhibited prominent LAMP2 and Ctsk staining within ruffled borders demarcated by F-actin rings (white arrows), the signal intensity of LAMP2 and Ctsk was drastically reduced in *Slc37a2*^KO^ osteoclasts, indicative of impaired ruffled border delivery. As a consequence, the ruffled borders of *Slc37a2*^KO^ osteoclasts appeared underdeveloped and immature when compared to the characteristic thin villous appearance of mature ruffled borders of WTs by TEM (**Fig. 7I**). Accordingly, ruffled border secretory activity was also diminished as indicated by the decreased levels of the cathepsin K cleavage product CTX-1(**Fig. 7J**), as well as a reduction in the levels of the active form of MMP9 (**Fig. 7K**) when media was sampled from cultures of bone-resorbing *Slc37a2*^KO^ and WT osteoclasts. Altogether, these data suggest that, during bone resorption, Slc37a2 facilitates the export of monosaccharide sugars from SLs that, in turn, prompts SL tubulation and transport to the ventral plasma membrane for secretion, and thus maintenance of the osteoclast ruffled border.

## DISCUSSION

Here we unmask the sugar transporter Slc37a2 as an obligatory component of osteoclast SLs and disclose several important findings that collectively advance our understanding of SL composition, organization and regulation. First, we provide a proteomic account of the osteoclast SL, thus expanding the molecular inventory of this enigmatic organelle. Second, we establish Slc37a2 as a *bona fide* constituent of SL membranes, assigning a home to an otherwise orphan transporter. Third, we reveal that SLs exist as highly dynamic tubular organelles in living osteoclasts that fuse with the ventral plasma membrane and service the ruffled border, an observation that has previously gone unnoticed. Fourth, we provide compelling genetic evidence that *Slc37a2* is essential for osteoclast function and bone metabolism, thus promoting Slc37a2 to the league of osteoclast transporters linked to the pathophysiological regulation of bone (Ribet et al., 2021). Finally, our work establishes Slc37a2 as a candidate SL monosaccharide exporter, addressing a long-standing question in the lysosome field.

Thus far, detailed studies of osteoclast SLs have been hampered by lack of suitable methods for isolating SLs to the scales and purity demanded for proteomic interrogation. Here, using SPIONs to exploit the endolysosomal pathway of osteoclasts, we have isolated and mapped the proteomic landscape of SLs for the first time. This unbiased approach unveiled a multitude of proteins enriched on SLs, including several proteins involved in membrane trafficking and molecular transport that have not been previously ascribed to SLs. Future work, including detailed biochemical and cellular characterization coupled with genetic models, will be required to validate independently the SL residency and pathophysiological contribution of these proteins to SL and bone homeostasis. Nevertheless, given that the SL proteome is enriched with genes linked to the homeostatic regulation of human bone mass, including those clinically-associated with monogenic skeletal dysplasia (e.g. *TCIRG1* and *CLCN7*) (Sobacchi et al., 2013), we anticipate that our organelle atlas will serve as a valuable resource to the community. The identification of Slc37a2 as a SL transporter associated with the regulation of bone mass in both humans (**Fig. 1E**) and in mice (**Fig. 3**) lends a proof-of-principle for this.

Originally identified as a cAMP-inducible gene in RAW264.7 macrophages (Takahashi et al., 2000), Slc37a2 is considered a member of the Slc37 subfamily (Slc37a1-a4) of transporters which reside in the endoplasmic reticulum (ER) and function as Pi-linked antiporters for sugar-phosphates such as G6P (Cappello et al., 2018). However, at least two lines of mechanistic and structural evidence hint that Slc37a2 is unlikely an ER-anchored G6P transporter. First, unlike the archetype Slc37 family member Slc37a4, which functions as a Pi-linked G6P antiporter coupled to G6Pases-a/b, the exchange activity of Slc37a2 is insensitive to the G6Pases inhibitor chlorogenic acid, indicating that Slc37a2 is not a physiological G6P transporter (Pan et al., 2011). Second, whereas Slc37a4 exists as a 10-transmembrane spanning protein localized to endoplasmic reticulum (ER) (Bartoloni and Antonarakis, 2004), Slc37a2 possess 12 transmembrane segments housing an extended N-terminal loop that undergoes extensive N-linked glycosylation (Kim et al., 2007) as well as localization signals (e.g. YxxØ) within its extreme C-terminus (**Fig. 2E**), criteria usually reserved for endolysosomal membrane proteins. Indeed, Slc37a2 has previously been detected on lysosomal membranes isolated from liver by proteomics (Chapel et al., 2013). In addition, during the course of this work, Slc37a2 was reported to localize to an endolysosomal-related organelle dubbed the ‘gastrosome’ in zebrafish microglia (Villani et al., 2019). Taken together with our extensive biochemical, morphological and functional studies, which localizes Slc37a2 to SLs, and/or late endosomes/lysosomes in non-osteoclastic cells (**Extended Data Fig.6**), we posit that the Slc37a2 transporter is a membrane resident of endolysosome-related organelles.

A remarkable feature of Slc37a2-bearing SLs is their extremely dynamic nature and ability to extend and maintain elongated tubules that fuse with the osteoclast bone-oriented plasma membrane. What is the role of ‘tubular’ SLs in osteoclasts during bone resorption? Based on our live cell results, we favor a model whereby SLs undergo microtubule-coupled extension and tubulation necessary to infiltrate the narrow infolds of the ruffled border for cargo delivery (**Fig. 8**). Although our ‘tubular SL’ model challenges the traditional paradigm of ruffled border genesis i.e. via the fusion of acidified ‘vesicles’ (Baron et al., 1985), it offers a simple yet plausible explanation to reconcile how SLs (typically 200-900 nm in diameter) (van Meel et al., 2011) are able to adapt to overcome the size constraints imposed by the tight membrane folds of the ruffled border (∼70-400 nm in diameter) (Segawa et al., 1989). This would also provide a mechanism for how collagenolytic enzymes, such as cathepsin K and MMP9, are continuously delivered to and secreted by the ruffled border during extended periods of bone resorption, such as occur during the formation of extended ‘trench-like’ resorptive trails (Borggaard et al., 2020, Merrild et al., 2015). Indeed, MMP9 secretion and activity has recently been shown to be mediated via tubular lysosomes in macrophages (Suresh et al., 2021). Alternatively, tubular SLs may serve to replenish components of the bone resorptive machinery that are lost from the ruffled border during membrane turnover, as occurs upon the engulfment of degraded bone matrix. The 116 kDa *a3* isoform (Tcirg1) of the V-ATPase proton pump, for example, is known to translocate from lysosomes to the bone-orientated osteoclast plasma membrane (Toyomura et al., 2003) and has been shown, in macrophages, to be transported via tubular lysosomes to phagosomes to establish the acidic environment hostile to pathogens (Sun-Wada et al., 2009). Given the analogies shared between the osteoclast ruffled border and phagolysosomal membranes, it is therefore conceivable that *a3* and/or other endolysosomal proteins (e.g. LAMP2) are recycled to the ruffled border via tubular SLs. Such a model would be in keeping with the: (i) impaired delivery of LAMP2 and cathepsin K to the ruffled border membrane; (ii) reduced extracellular secretion of cathepsin K and active-MMP9; (iii) altered ruffled border morphology and; thus (iv) resorptive deficits exhibited by *Slc37a2*^KO^ osteoclasts.

**Fig. 8.**
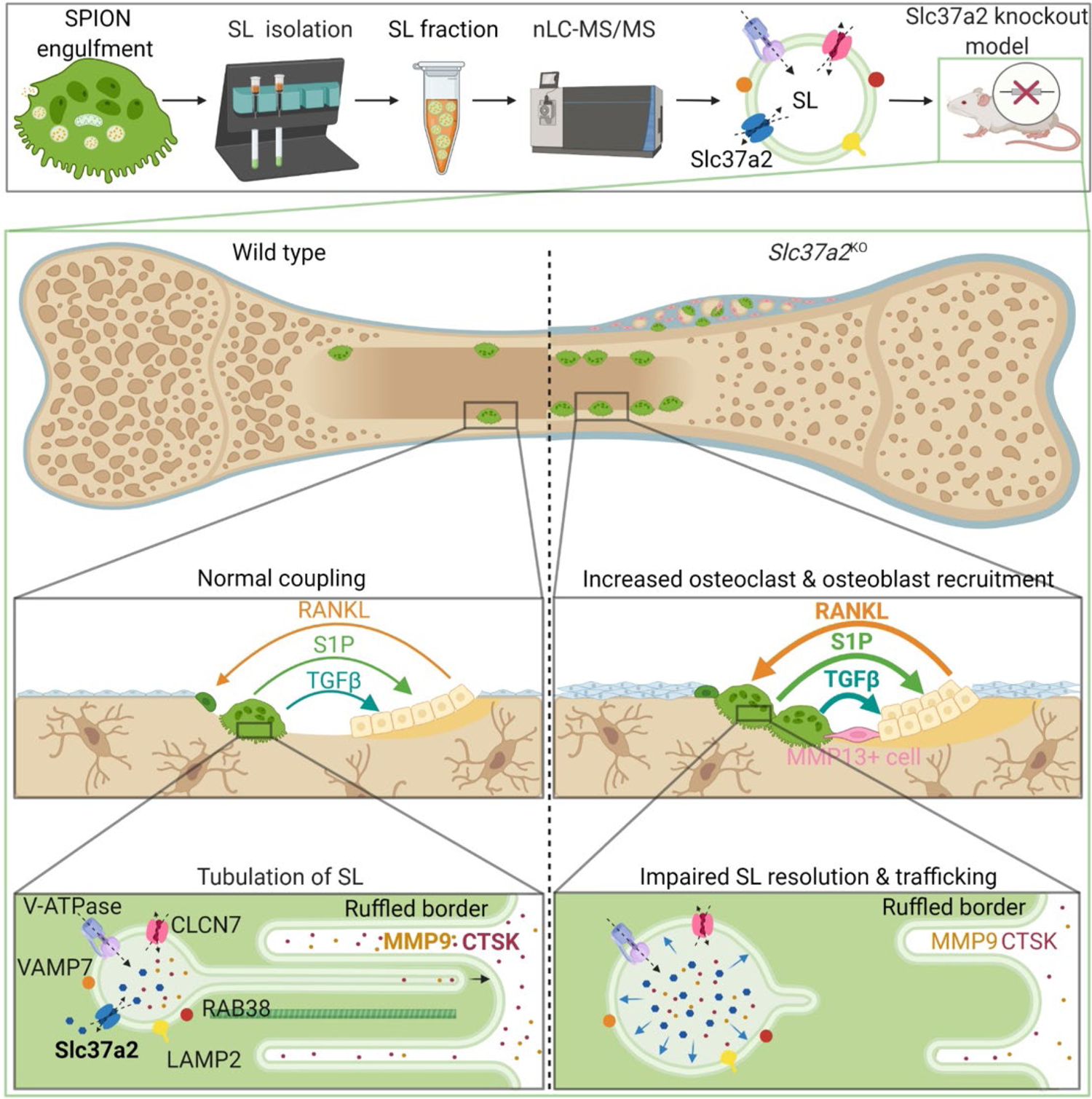
Working model of Slc37a2 in osteoclast function and bone metabolism. Top panel provides an overview of proteomic identification of sugar transporter Slc37a2 on osteoclast secretory lysosomes, middle panels highlights tissue level alterations in osteoclast-osteoblast coupling during bone remodeling in Slc37a2-deficient mice and bottom panel illustrates the proposed role for Slc37a2 in the regulation of sugar export from secretory lysosomes, a prerequisite for secretory lysosome resolution, membrane tubulation and delivery to the osteoclast ruffled border.

Perhaps the most striking observation of this study is the dramatic high bone mass phenotype in *Slc37a2*^KO^ mice. This phenotype is attributed primarily to impaired bone resorption by osteoclasts leading to imbalanced remodeling-based bone formation by osteoblasts. *In vivo*, this deficit in osteoclast function appears partially compensated for by enhanced osteoclastogenesis via increased RANKL/OPG levels, as well as increased expression of key proteases known to participate in collagen-matrix digestion (e.g. MMP9, MMP13, MMP14) (Zhu et al., 2020) and osteoclast/matrix-derived coupling factors that promote remodeling-based bone deposition by osteoblasts (e.g. Sphk1/S1P and TGF-β) (Sims and Martin, 2020). Altogether, these phenotypic and cellular features are strikingly reminiscent of those reported for cathepsin K-deficient mice (Saftig et al., 1998, Lotinun et al., 2013, Kiviranta et al., 2005). What distinguishes *Slc37a2*^KO^ mice from mice lacking cathepsin K, and other models of osteosclerosis attributed to osteoclast dysfunction e.g. β3-integrin (McHugh et al., 2000) and Plekhm1 (Van Wesenbeeck et al., 2007), is the conspicuous fibro-cartilaginous lesion observed within the physeal-metaphyseal interface (**Fig. 4**). While the pathoetiology of this lesion requires further study, it may reflect failure of *Slc37a2*^KO^ osteoclasts to complete bone ‘cut-back’ i.e. to decrease bone width within the periosteal metaphyseal area during longitudinal growth (Orwoll, 2003). Alternatively, it may point to disturbances in a population of matrix-degrading cells specific to the periosteal-bone niche that are distinct from osteoclasts but are functionally dependent on Slc37a2. For example, septoclasts, a rare but specialized mesenchymal stromal-derived cell type, have recently been shown to resorb cartilage during endochondral ossification and fracture healing and express high levels of MMP9, 13 and 14 (Sivaraj et al., 2022), in keeping with the increased MMP9/13/14 expression within *Slc37a2*^KO^ femurs observed here (**Fig.s 4H, 5K&P**). Whether septoclasts, reversal cells, or an as yet undefined Runx2/MMP13^+^ matrix-degrading periosteal cell type contribute directly to the *Slc37a2*^KO^ bone phenotype requires the future generation and detailed characterization of several cell-specific conditional *Slc37a2* knockout mouse lines, which forms the basis of our ongoing investigations.

Previous biochemical studies have described the transport of glucose and fructose out of lysosomes by unknown monosaccharide transporters (Mancini et al., 1990, Lizak et al., 2019). Our results suggest that, in osteoclasts, Slc37a2 fulfils this role. First, metabolic profiling demonstrated that loss of Slc37a2 led to increased levels of glucose and fructose in osteoclasts. Second, the morphological swelling of SLs in *Slc37a2*^KO^ osteoclasts together with the impaired capacity of ‘sucrosomes’ to undergo invertase-induced resolution, indicates that the export of glucose and fructose from SLs is inhibited when Slc37a2 is lacking, in keeping with a lysosomal storage disorder. Considering that the extrusion of monosaccharides is an established prerequisite for lysosomal resolution (Swanson et al., 1986), tubulation (Bright et al., 2016) and transport (Bandyopadhyay et al., 2014), we envisage a membrane remodeling cascade whereby Slc37a2-mediated export of glucose and fructose drives the osmotic extrusion of water from SLs, which in turn, reduces their hydrostatic tension, thus prompting SL tubulation and transport to the ruffled border (**Fig. 8**).

In recent years, the critical importance of SLCs in human metabolic diseases and their largely untapped potential as ‘druggable’ targets has gained increasing momentum (Schumann et al., 2020, Lin et al., 2015, Superti-Furga et al., 2020). Altogether, our findings that *Slc37a2* deletion increases skeletal bone mass and strength and attenuates age-associated bone loss, lends a rationale for the development of Slc37a2 inhibitors as a potential therapeutic treatment for metabolic bone diseases like osteoporosis.

## ACKNOWLEDGEMENTS

We thank Rob Day and Alex Haynes (Royal Perth Hospital, W.A.) for their technical assistance with the biomechanical testing studies, Lisa Griffiths (PathWest, W.A.) for performing electron microscopy on bone tissue, Daniel Yagoub for proteomics assistance, James Rae for assistance with electron microscopy of cultured cells and Natalie Sims (St Vincent’s Institute, V.I.C) for insightful discussions. We are also indebted to Michael S. Marks (University of Pennsylvania, U.S.A.) and Allistair N. Hume (The University of Nottingham, U.K.) and Roland Baron (Harvard School of Dental Medicine, U.S.A.) for the provision of plasmids. This work was supported by NHMRC Project funding APP1143921 to N.J.P., R.D. and J.X., NHMRC grants APP1140064 and APP1150083 and fellowship APP1156489 to R.G.P., a NHMRC grant APP2003629 and Department of Health Western Australia Merit Award 1186046 to B.M., an Arthritis Australia and HJ & GJ Mackenzie Grant (N.J.P and P.Y.N), and a Faculty of Health and Medical Sciences Research Grant Scheme (SE Ohman Medical Research Fund to N.J.P. and P.Y.N). A.B.P.R is supported by Australian Government Research Training Program Scholarship. The authors acknowledge the facilities, and the scientific and technical assistance of Microscopy Australia at the Centre for Microscopy, Characterisation & Analysis, The University of Western Australia, a facility funded by the University, State and Commonwealth Government and of the Microscopy Australia Research Facility at the Centre for Microscopy and Microanalysis at The University of Queensland. N.J.P. and K.S. are supported by COST Action GEMSTONE CA18139 (European Cooperation in Science and Technology).

## CONTRIBUTIONS

P.Y.N. performed the experiments, analyzed the data contributed to manuscript drafting and editing. A.B.P.R. performed experiments, analyzed data contributed to manuscript drafting and figures, G.Q. performed micro-CT analyses, cryosectioning and biomechanical testing. B.H.M. performed GWAS analysis and contributed figures, J.W.Y.T., contributed *in vitro* experiments and metabolomics, E.L.B., performed micro-CT and histomorphometric analyses, S.S. contributed to the establishment of microinjection experiments, K.C., assisted with micro-CT analyses and OVX-studies, L.A. assisted with proteomic analyses, M.B. contributed to the metabolomic analyses, E.T.T.T.N. assisted with histomorphometric analyses, J.K. analyzed the RNAseq data, J.M.P. performed histopathological analyzes and critically reviewed the manuscript, K.S. analyzed the *in vitro* bone resorption data and edited and critically reviewed the manuscript, R.D.T. and J.X. financially supported and critically reviewed the manuscript, R.G.P. performed APEX work, edited and critically reviewed the manuscript, H.T. offered critical suggestions to the project design and manuscript drafting, N.J.P. conceptualized the project, designed all experiments, wrote the manuscript and financially supported the project.

## METHODS

### Generation of Slc37a2 antibodies

A Slc37a2-specific polyclonal antibody was generated by immunizing rabbits with the peptide ‘CTPPRHHDDPEKEQ,’ corresponding to the cytoplasmic facing intracellular loop of mouse Slc37a2. Antisera were affinity-purified against the corresponding immunization-peptide. Slc37a2 peptide antibodies were produced using the PolyExpress^TM^ service by GenScript (New Jersey, USA).

### Additional antibodies

#### Primary

Rabbit monoclonal Akt (pan) (C67E7), (Cell Signaling, Cat#: 4691)

Rabbit monoclonal Phospho-Akt (Ser473) (D9E) XP®, (Cell Signaling, Cat#: 4060)

Mouse monoclonal anti-Cathepsin K, (clone 182-12G5, Millipore, Cat#: MAB3324)

Mouse monoclonal anti-Cathepsin K (E-7), (Santa Cruz, Cat#: sc-48353)

Rabbit polyclonal anti-Cathepsin B (Atlas Antibodies, Cat#: HPA018156)

Rabbit polyclonal anti-Collagen I antibody, (Abcam, Cat#: ab34710)

Rabbit polyclonal anti ERK1/2, (Promega, Cat#: V1141)

Mouse monoclonal anti-actin deposited by Lin, J.J.-C, (DSHB, Cat#: JLA20)

Rat monoclonal anti-LAMP-2 deposited by August, J.T, (DSHB, Cat#: ABL-93)

Rat monoclonal anti-LAMP-2 deposited by Granger, B. L, (DSHB, Cat#: GL2A7)

Mouse monoclonal anti-NFATc1 deposited by Crabtree, G.R, (DSHB, Cat#: 7A6)

Rabbit polyclonal anti-MMP13 (ab39012), (Abcam, Cat#: ab39012)

Rabbit polyclonocal anti-Rab1b (Santa Cruz, Cat#: sc-599)

Mouse monoclonal anti-Rab5, (Synaptic Systems, Cat#: 108 011)

Rabbit monoclonal anti-Rab7 (D95F2), (Cell Signaling, Cat#: 9367)

Mouse monoclonal anti-Rab38 (A-8), (Santa Cruz, Cat#: sc-390176)

Rabbit monoclonal anti-RUNX2 (D1L7F), (Cell Signaling, Cat#: 12556)

Rabbit polyclonal anti-TCIRG1, (Abcam, Cat#: ab139812)

Rabbit polyclonal anti-VGLUT-1, (Synaptic Systems, Cat#: 135 303)

#### Secondary

Goat anti-mouse IgG (Fab specific)-Peroxidase antibody, (Sigma-Aldrich, Cat#: A9917)

Goat anti-rabbit IgG (whole molecule)-Peroxidase antibody, (Sigma-Aldrich, Cat#: A0545)

Alexa Fluor 488 goat anti-mouse IgG (H+L), highly cross-adsorbed, (Thermo Fisher Scientific, Cat#: A-11029)

Alexa Fluor 568 goat anti-mouse IgG (H+L), highly cross-adsorbed, (Thermo Fisher Scientific, Cat#: A-11031)

Alexa Fluor 647 goat anti-mouse IgG (H+L), highly cross-adsorbed, (Thermo Fisher Scientific, Cat#: A-21236)

Alexa Fluor 488 goat anti-rabbit IgG (H+L), highly cross-adsorbed, (Thermo Fisher Scientific, Cat#: A-11034)

Alexa Fluor 568 goat anti-rabbit IgG (H+L), highly cross-adsorbed, (Thermo Fisher Scientific, Cat#: A-11036)

Alexa Fluor 647 goat anti-rabbit IgG (H+L), highly cross-adsorbed, (Thermo Fisher Scientific, Cat#: A-21245)

Alexa Fluor 555 donkey anti-rat, IgG (H+L), highly cross-adsorbed, (Thermo Fischer, Cat#: A48270)

### Animals

Slc37a2 gene-trapped ES cells (JM8A3.N1 cell line, Clone ID EPD0641_4_E06) were obtained from the KOMP consortium and provided by the UC Davis KOMP Repository. Slc37a2 knockout (KO) mice were generated through the Australian Phenomics ES Cell-to-Mouse Service (Monash University). All mice were maintained on the C57BL/6N background under specific-pathogen free conditions at the Animal Resources Centre (ARC), Murdoch, Western Australia (W.A). Mice were maintained in a temperature and humidity-controlled room on a 12 h light cycle with *ad libitum* access to water and a standard laboratory chow diet. The study was performed in strict accordance with the Animal Welfare Act 2002 (W.A) and requirements of the eighth (2013) edition of the Australian code for the care and use of animals for scientific purposes. All the animals were handled according to institutional animal care protocols approved by the Animal Ethics Committee of The University of Western Australia (Approval No. RA13/100/1475).

### Genotyping

The genotype of mice was determined by PCR using the cycling parameters recommended by UC Davis KOMP Repository. The presence of Slc37a2 WT and tm2a alleles in the mice genome were analyzed by multiplex PCR using primer sequences CSD-loxF: GAGATGGCGCAACGCAATTAATG, CSD-R: GACCTAGCTTCTCAGGCATCTCAGG, CSD-Forward: AGAAGGGGTACAGAAGCAACAGC, CSD-ttReverse: CCATGTGTCCCTTCATGCTTTAGG.

### Bone marrow macrophage extraction and osteoclast culture

Bone marrow macrophages (BMMs) were harvested from the long bones of Slc37a2^+/+^ wild-type (WT) and Slc37a2^-/-^ (KO) mice in accordance with the UWA Institutional Animal Ethics Committee Guidelines. Unless otherwise specified, all cell culture reagents were purchased from Thermo Fisher Scientific. Briefly, extracted primary cells were cultured in complete α-MEM containing 10% fetal bovine serum (FBS), 2mM L-glutamine, 50U/ml penicillin, 50U/ml streptomycin, and supplemented with M-CSF (25ng/ml). All cultures were maintained in humidified conditions of 5% CO2 at 37°C. Osteoclasts were generated from BMMs differentiated with RANKL (10ng/ml, R&D Systems) and M-CSF (2ng/ml) *in vitro*. Cells were then harvested and processed for RNA extraction, immunoblot or proteomic analysis, or seeded onto glass coverslips, glass-bottomed culture dishes (ibidi), cortical bone slices (Boneslices.com) or Osteo Assay Surface Stripwells (Corning).

### Secretory lysosome extraction

Secretory lysosome (SL) extraction was adapted using methods described previously (Nakamura et al., 2014, Walker and Lloyd-Evans, 2015), but with the following modifications. Briefly, a large scale osteoclastogenesis was performed on 4 x 15cm diameter dishes at a cell density of 5 × 10^6^ BMMs per plate. Following osteoclast formation, the cells were ‘pulsed’ for 24h with 10µl of superparamagnetic iron oxide nanoparticles (SPIONs) (carboxyl-Adembeads; Ademtech) per dish and ‘chased’ for an additional 24h to accumulate SPIONS in secretory lysosomes. Cells were washed with warm PBS and scraped in homogenisation buffer (10mM HEPES-KOH pH 7.4, 250mM sucrose, 5mM DTT, 1mM EDTA, and protease inhibitor cocktail, Roche). The cells were then homogenized using an Eberbach Corporation (E2355) Con-Torque Tissue Homogenizer at the highest rpm setting, a PYREX® 5mL Potter-Elvehjem and Pestle (7725T-5) for 30 strokes then passed 8 times through a 23G needle. Lysates were collected and centrifuged at 200 × g for 10min at 4°C to remove debris and unbroken cells. The supernatants were then loaded onto a Miltenyi Biotec MS Column (130-042-201) equilibrated with 1ml 0.5% BSA in PBS and immobilized SPIONs washed with DNase solution (1mg/ml DNase, 0.1mM sucrose in PBS). Fractions (F1-F3) containing enriched SLs were collected by separating the column from the Miltenyi Biotec MiniMACS™ Separator magnet and sequentially eluting the column with 200µl of elution buffer (0.1mM sucrose, protease inhibitor cocktail in PBS). Aliquots of the three enriched SL fractions (F1, F2, F3), the cell homogenate (H), the post-nuclear supernatant (PNS) and the column flow-through (UB) were snap frozen in liquid N_2_ and stored at −80°C until required. Successful isolation of SLs was confirmed using immunoblotting for specific organelle markers, transmission electron microscopy and by an enzymatic assay for ß-Hexosaminidase activity.

### Proteomic sample preparation

Protein concentration of fractions was determined by Bradford assay (Bio-Rad #5000006). 10 µg of the total cell homogenates and SL fraction F2 (each corresponding to a biological triplicate) were used for proteomic analysis. Briefly, samples were mixed with SDS-PAGE sample buffer and denatured before 1-dimensional electrophoresis (1-DE) through an Invitrogen precast 10% NuPAGE Bis-Tris gel with corresponding NuPAGE running buffer at 80V for 1h. The gel was washed with Milli-Q H2O and fixed with 40% ethanol, 10% acetic acid in Milli-Q H_2_O for 15min with gentle rocking. After a series of Milli-Q H_2_O washes, the gel was stained using a pre-prepared Bio-Rad Colloidal Coomassie stain (#1610803) and then destained with Milli-Q H_2_O. The gel lanes were then divided into ten fractions. The ten gel fractions were further divided into 1mm square cubes, further de-stained and then digested in-gel with trypsin to extract peptides according to the methods of (Shevchenko et al., 2006). Peptides were dried using a Speed-Vac and analyzed by mass spectrometry.

### LC-MS/MS analysis

For mass spectrometry analysis, peptides were separated at a flow rate of 300nl/min on a 50 cm long, 75µm internal diameter EASY-spray PepMap C18 column (Thermo Fisher Scientific) using a Dionex UltiMate 3000 Nano-UHPLC system (Thermo Fisher Scientific). The column was maintained at 50°C. Buffer A and B were 0.1% formic acid in water and 0.1% formic acid in acetonitrile respectively. Peptides were eluted in 165min runs, with a gradient from 3-5% buffer B for 5min, from 5-25% buffer B for 105min, from 25-35% buffer B for 15min, from 35-95% buffer B for 20min and equilibration (3% buffer B) for 19min. Eluting peptides were analyzed on an Orbitrap Fusion mass spectrometer (Thermo Fisher Scientific). The instrument was operated in a data dependent, “Top Speed” mode, with cycle times of 3sec. Peptide precursor mass to charge ratio (m/z) measurements (MS1) were carried out at 120000 resolution in the 375 to 1500 m/z range. The MS1 AGC target was set to 4e5 and maximum injection time to 50msec. Precursor priority was set to “most intense” and precursors with charge state 2 to 7 only were selected for HCD fragmentation. Fragmentation was carried out using 35% collision energy. The m/z of the peptide fragments were measured in the ion trap using an AGC target of 1e4 and 50msec maximum injection time.

### Data analysis

Mass spectrometry spectral data were analyzed using Proteome Discoverer™ 2.3 Software (Thermo Fisher Scientific). Briefly, the reference FASTA library used for analysis was downloaded from UniProtKB (5/02/2020) for *mus musculus* containing 86,447 reviewed and un-reviewed peptides. The cleavage sites were set to correspond to Trypsin and maximum missed cleavage sites was set at 2. Minimum and maximum peptide lengths were set at 7 and 144 respectively with the precursor mass tolerance set at 4.4ppm. The biological triplicate peptide abundances were pooled and abundance ratios were calculated for total cell homogenate pooled triplicates over SL (F2) pooled triplicates.

### Analysis of human orthologues

Enrichment of genome-wide association study (GWAS) associations among human homologues of the mouse SL gene set was performed using the VEGAS2Pathway function of the VErsatile Gene-based Association Study 2 (VEGAS2) software package (Mishra and MacGregor, 2017). The VEGAS2Pathway approach tests for enrichment of significant associations in GWAS summary data in pre-defined gene sets, while correcting for gene lengths, linkage disequilibrium (LD) between markers and the size of the gene set. Gene analysis windows included 50Kb on either side of each gene, with the 1000 Genomes phase 3 EUR dataset used as the LD reference. A custom gene set was created for the analysis consisting of the human homologues of the top 218 enriched mouse genes identified in our studies.

### Sample preparation for cryosectioning and immunohistochemistry

Hindlimbs of mice were dissected and muscles were thoroughly removed before the femurs were fixed in 10% formalin at 4°C overnight. Samples were washed thrice with PBS before decalcification in cold, 14% EDTA pH 8.0 for 48h. Decalcified bone samples were incubated in 15% and 30% sucrose (Sigma-Aldrich) in PBS each for 48h at 4°C, then embedded in SCEM-L1 embedding medium (SECTION-LAB), and rapidly cooled using a hexane-dry ice coolant mix. Bone cryosections (20-30µm) were generated with a tungsten carbide blade (ProSciTech) and Leica CM1900 cryostat, using Kawamoto’s film method (Kawamoto and Shimizu, 2000).

For immunostaining, cryosections were completely thawed at RT and rehydrated with PBS at RT for 5min to remove the embedding medium. Sections were permeabilized with 0.3% Triton-X/PBS at RT for 20min and blocked with 5% BSA/PBS at RT for 30min. Primary antibodies were incubated at 4°C overnight. Primary antibodies included: Slc37a2 (rabbit polyclonal, this paper, 1/100 dilution), RUNX2 (rabbit monoclonal, Cell Signaling, 1/200 dilution), MMP13 (rabbit polyclonal, Abcam, ab39012 1/300 dilution). Conjugate antibodies and fluorescent probes included: Rhodamine phalloidin (Cat. No. R415, 1/500 dilution), Alexa Fluor 647 Phalloidin (Cat. No. A22287, 1/300 dilution), IVISense Osteo 680 (PerkinElmer, 1/300 dilution), IVISense Integrin Receptor (IntegriSense^TM^) 645 (PerkinElmer, 1/300 dilution). Following primary antibody incubation, sections were washed and incubated with secondary antibodies at RT for 2h. Secondary antibodies used included: anti-rabbit Alexa Fluor 488 (Thermo Fisher, 1/500 dilution) and nuclei were stained using Hoechst 33258 (Cat. No. H3596, Invitrogen, 2µg/ml). Sections were washed, mounted onto glass slides using ProLong Glass antifade mountant (Cat. No. P36980, Invitrogen) and sealed with glass coverslips and imaged by confocal microscopy. Sections were imaged using the NIKON A1R in a Ti-E inverted motorized confocal microscope equipped with Nikon PlanApo 20x (air), Nikon PlanApo 60x WI IR, 1.27NA (oil) and Nikon PlanApo 100x, 1.45 NA (oil) immersion lenses.

### Protein extraction and immunoblotting

Cells were lysed on ice in RIPA buffer supplemented with Protease Inhibitor Cocktail (Cat#. 11873580001, Roche). Lysates were cleared by centrifuging at 15,000g and 4°C for 20min (Sorvall Legend Micro 17R Centrifuge, Cat#. 75002440, Thermo Fisher Scientific). Protein concentrations were determined through Bradford assays (Cat#. 5000006, Bio-Rad). For immunoblotting, loading buffer was added and samples incubated at 37°C for 1h before SDS-PAGE electrophoresis and transfer onto nitrocellulose membranes. Membranes were blocked with 5% skim milk in TBST, and primary antibodies incubated overnight at 4°C in TBST with 1% skim milk. Primary antibodies include: LAMP-2 (DSHB ABL-93, 1/1000 dilution), Rab7 (Cell Signaling 95746, 1/2000 dilution), Ctsk (Santa Cruz sc-48353, 1/250 dilution), Rab38 (Santa Cruz sc-390176, 1/200 dilution), Rab5 (Synaptic Systems 108 011, 1/1000 dilution), Stx16b (Synaptic Systems 110 163, 1/300 dilution), Rab1b (Santa Cruz sc-599, 1/5000 dilution), VDAC3 (Santa Cruz sc-292328, 1/500 dilution), VGluT1 (Synaptic Systems 135 303, 1/1000 dilution), Slc37a2 (this paper, 1/1000 dilution), V-ATPase D1 (Santa Cruz sc-81887, 1/1000 dilution), NFATc1 (DSHB, 7A6 1/1000 dilution), c-Src (Millipore Merck: 05-184, clone GD11, 1/1000 dilution), Ctsb (Atlas Antibodies, Sigma Aldrich HPA018156, 1/250 dilution), pERK1/2 (Santa Cruz sc-7383, 1/500 dilution), t-ERK1/2 (Promega V114A, 1/1000 dilution), p-AKT (Cell Signaling Technology, 1/1000 dilution), t-AKT (Cell Signaling Technology, 1/1000 dilution), IκBα (Santa Cruz: sc-371, 1/1000 dilution) and β-actin (DSHB JLA20, 1/2000 dilution). Membranes were next washed and incubated with peroxidase-conjugated secondary antibodies (1:5000) in TBST with 1% skim milk for 1h. Membranes were developed with Western Lightning Ultra Chemiluminescence reagent (Cat#: NEL113001EA, Perkin Elmer), and imaged with an ImageQuant LAS 4000 (GE Healthcare).

### RNA extraction and quantitative PCR

Bone tissue samples were harvested from mice, snap frozen in liquid nitrogen, and ground into powder using a hammer. The powdered bone tissue samples were transferred into RNAse-free microcentrifuge tubes and RNA was extracted using TRIzol reagent (Cat#. 15596026, Invitrogen), followed by purification using the PureLink RNA mini kit (Cat#. 12183020, Invitrogen). For cell cultures, media was removed before addition of TRIzol reagent. Following extraction, RNA samples were treated with RNase-free DNase I (Cat#. 18068015, Invitrogen). For first-strand synthesis of cDNA, 1-2µg of total RNA was reverse transcribed using SensiFAST^TM^ cDNA Synthesis Kit (Cat#. BIO-65054, Meridian Bioscience) according to manufacturer’s protocol. Quantitative PCR was performed using SensiMix^TM^ II Probe Kit (Cat#. BIO-83005) with Universal ProbeLibrary (Roche Diagnostics), and the CFX96 Touch System (Bio-Rad). Relative fold-expression was normalized to β-actin (ACTB; TaqMan), hydroxymethylbilane synthase (HMBS) and hypoxanthine-guanine phosphoribosyltransferase (HPRT) housekeeping controls using primer sequences listed below.

### Oligonucleotides for qPCR

Primer: SLC37A2 isoform 2 Forward: tgctgcatcatgctgatctt; Reverse: cccattctggccaatgtagt

Primer: RUNX2 Forward: cgtgtcagcaaagcttctttt; Reverse: ggctcacgtcgctcatct

Primer: MMP9 Forward: agacgacatagacggcatcc; Reverse: tcggctgtggttcagttgt

Primer: MMP13 Forward: cagtctccgaggagaaactatgat; Reverse: ggactttgtcaaaaagagctcag;

Primer: MMP14 Forward: gagaacttcgtgttgcctga, Reverse: ctttgtgggtgaccctgact

Primer: WNT16 Forward: catgaatctacacaacaacgaagc, Reverse: ttttccagcaggttttcaca;

Primer: WNT10b Forward: aatgcggatccacaacaac, Reverse: ctccaacaggtcttgaattgg

Primer: TGFβ1 Forward: tggagcaacatgtggaactc, Reverse: gtcagcagccggttacca

Primer: HPRT Forward: ggagcggtagcacctcct, Reverse: ctggttcatcatcgctaatcac

Primer: ACP5 Forward: cgtctctgcacagattgcat, Reverse: aagcgcaaacggtagtaagg

Primer: CTSK Forward: cgaaaagagcctagcgaaca, Reverse: tgggtagcagcagaaacttg

Primer: HMBS Forward: cagtgatgaaagatgggcaac, Reverse: aacagggacctggatggtg

Primer: ALP Forward: cggatcctgaccaaaaacc, Reverse: tcatgatgtccgtggtcaat

Primer: SOST Forward: atcccagggcttggagagta, Reverse: ccggttcatggtctggtt

Primer: SPHK1 Forward: tgtgaaccactatgctgggta, Reverse: gcagcccagaagcagtgt

Primer: SEMA4D Forward: aagtgggtgcgctacaatg, Reverse: gggcctcactgtcgatacac

Primer: BMP6 Forward: acatggtcatgagctttgtga, Reverse: gtcgttgatgtggggagaac

Primer: RANK Forward: gtgctgctcgttccactg, Reverse: agatgctcataatgcctctcct

Primer: RANKL Forward: tgaagacacactacctgactcctg, Reverse: cccacaatgtgttgcagttc

Primer: OPG Forward: gtttcccgaggaccacaat, Reverse: ccattcaatgatgtccaggag

### Immunofluorescence confocal microscopy

To preserve the integrity of the tubular SL network, rapid cellular fixation under precise temperature control and microtubule stabilization was performed according to the methods described in (McGrath et al., 2021). Briefly, osteoclasts were placed on a 37°C heating block prior to the addition of freshly made, pre-warmed (37°C) 8% PFA in 2x microtubule stabilizing buffer (MTSB, 160mM PIPES pH 6.8, 10mM EGTA and 2mM MgCl_2_) directly to the cell culture media in a 1:1 ratio (final concentration of 4% PFA in 1x MTSB). The cells were then immediately returned to a 37°C incubator for 15min. Following fixation, cells were rinsed gently thrice in pre-warmed PBS to remove residual fixative and stored in PBS at 4°C until required.

For immunostaining, osteoclasts were permeabilized with 1% saponin in PBS for 5min at RT, and then washed twice with PBS. Cells were blocked in 3% BSA/PBS for 30min at RT. Cells were incubated with a rabbit polyclonal antibody specific to mouse Slc37a2 (this study, 1/200 dilution) and/or a rat monoclonal LAMP-2 antibody (GL2A7, DSHB, 1/100 dilution) in 0.2% BSA/PBS containing 0.1% saponin overnight at 4°C. Cells were washed four times with 0.2% BSA/PBS and incubated with donkey anti-rabbit Alexa Fluor-488 (Thermo Fisher Scientific, 1/500 dilution), donkey anti-rat Alexa Fluor-555 (Thermo Fisher Scientific, 1/500 dilution) and Hoechst 33258 (Thermo Fisher Scientific, 1/10,000 dilution) diluted in 0.2% BSA/PBS containing 0.1% saponin for 1h at RT. Finally, cells were washed thrice with PBS and samples mounted onto glass slides using ProLong Glass Antifade mountant (Invitrogen). Cells were imaged using the NIKON A1R in a Ti-E inverted motorized confocal microscope equipped with Nikon PlanApo 20x (air), Nikon PlanApo 60x WI IR, 1.27NA (oil) and Nikon PlanApo 100x, 1.45 NA (oil) immersion lenses.

### Microinjection

Primary BMMs were cultured directly onto ibidi 35mm glass bottom culture dishes (Cat# 81218-200, ibidi) under osteoclastogenic condition (10ng/ml RANKL and 25ng/ml M-CSF) for a period of 4 days. Nuclear microinjection of plasmid vectors including ^mCherry-^Slc37a2 isoform 1, ^emGFP-^Slc37a2 isoform 2, ^RFP-^LAMP1, Cortactin^-dtTomato^, ^mCherry-^VAMP7, ^mCherry-^Rab38) and ^mCherry-^Rab5 (all 1µg/µl in water) was performed on a Olympus IMT-2 inverted microscope, using Femtotip II microinjection capillaries (Cat#. 5242957.000, Eppendorf) combined with the Eppendorf electronic microinjector FemtoJet 4x (Cat#. 5253000033, Eppendorf) and InjectMan®4 micromanipulator (Cat#. 5192000035, Eppendorf) using external continuous pressure. After microinjection, cell media was refreshed and the osteoclasts were maintained at 37°C and 5% CO_2_ overnight before live cellular imaging the following day.

### Live cell imaging

Primary BMMs were cultured directly onto ibidi 35mm glass bottom culture dishes under osteoclastogenic conditions (10ng/ml RANKL and 25ng/ml M-CSF) for a period of 4-5 days. For live imaging of osteoclasts on bone, similar conditions as above were employed to culture primary BMMs directly on bovine cortical bone slices (Boneslices.com) for 8-10 days, before the bone slices were transferred to a glass bottom culture dish for imaging. Prior to imaging, osteoclasts were incubated with live imaging compatible fluorescent probes: SIR-Actin (Cytoskeleton, Inc.), LysoTracker^TM^ Red DND-99 (Thermo Fisher), LysoTracker^TM^ Green DND-26 (Thermo Fisher), DQ^TM^ Red BSA (Thermo Fisher), OsteoSense 680 fluorescent probe (Perkin Elmer), Wheat Germ Agglutinin (Thermo Fisher), Alexa FluorTM 488 conjugate (Thermo Fisher), Magic Red Cathepsin K Assay (ImmunoChemistry Technologies LLC), MitoTrackerTM Red CMXRos (Thermo Fisher), Hoechst 33258 Pentahydrate (bis-Benzimide) (Thermo Fisher), according to manufacturer’s instructions.

Live osteoclasts treated with fluorescent live probes or microinjected with plasmids of interest were imaged using the NIKON A1R in a Ti-E inverted motorized confocal under controlled atmospheric conditions (37°C and 5% CO_2_) using an Oko Labs stage top incubator. Frames were captured at indicated time-points using NIS Elements software (Nikon). Images were processed and channels pseudo-colored using ImageJ (Fiji) software (Schindelin et al., 2012). For sucrose-mediated enlargement of lysosomes, BMM-derived osteoclasts from WT and *Slc37a2*^KO^ mice were incubated with 30mM sucrose in complete α-MEM for 24h after which cells were incubated with LysoTracker^TM^ Red DND-99 dye for 30 min to label sucrosomes. Sucrosome resolution was initiated by the addition of 0.5 mg/ml, of invertase (*S. cerevisiae*) for 1h and then images captured continuously by time-lapse confocal microscopy for a period of 10min.

### Constructs

^mCherry-^Slc37a2 iso1 and ^emGFP-^Slc37a2 iso2 were generated using Genscript Express Cloning SC1691 cloning service. Briefly, ^mCherry-^Slc37a2 iso1 was designed by cloning the full length coding sequence (CDS) of Slc37a2 variant 1 (NCBI reference sequence: NM_001145960.1) into the pcDNA3.1+ vector. The initial ATG start codon was removed and replaced with a mCherry sequence, followed by a 21bp linker sequence (5’-TCCGGACTCAGATCTCGAGCG-3’) downstream. ^emGFP-^Slc37a2 iso2 was generated by cloning the full length CDS of Slc37a2 variant 2 (NCBI reference sequence: NM_020258.4) into the pcDNA3.1+ vector. Similarly, the initial ATG start codon was removed and replaced with an emGFP sequence, followed by a 21bp linker sequence (5’-TCCGGA CTCAGATCTCGAGCG-3’) downstream.

To create the entry clone required for gateway cloning into the lentiviral destination vector, SalI and NotI sites were inserted into ^emGFP-^Slc37a2 iso2 using the following primers (5’-TAAGCAGTCGACATGGTGAGCAAGGGCGAG-3’; 5’-TGCTTAGCGGCCGCTCAAATTTGTTTGTACCCACTGC-3’), then ligated into the pENTR1A no ccDB (w48-1) vector (a gift from Eric Campeau and Paul Kaufman, Addgene plasmid # 17398; http://n2t.net/addgene:17398; RRID:Addgene_17398) using the Quick Ligation^TM^ kit (New England BioLabs) according to manufacturer’s instructions. The pLenti PGK Hygro emGFP-Slc37a2 iso2 vector (referred to as pLenti ^emGFP-^Slc37a2) was constructed by recombining pENTR1a emGFP-Slc37a2 iso 2 with the pLenti PGK Hygro DEST (w530-1) vector (a gift from Eric Campeau and Paul Kaufman, Addgene plasmid # 19066; http://n2t.net/addgene:19066; RRID:Addgene_19066) using Gateway LR Clonase II enzyme mix (Thermo Fisher) as per manufacturer’s instructions.

### Lentiviral production and transduction of primary bone macrophages

To produce lentivirus, pLenti ^emGFP-^Slc37a2 and pHIV-PV-VSVG (kind gift from Dr. Scott A. Fisher, The University of Western Australia) packaging plasmid were co-transfected into HEK293FT cells (Thermo Fisher) using TransIT-Lenti transfection reagent (Mirus) according to manufacturer’s instructions. 18h post transfection, media containing the transfection reagent was removed and replaced with fresh, high-BSA DMEM (Gibco) containing 10% FBS (Gibco) and 0.1g/L bovine serum albumin (Sigma-Aldrich). Culture media containing lentivirus were collected 48-72h post transfection and concentrated using lentivirus concentration solution (Origene) according to manufacturer’s instructions. The lentiviral pellet obtained post concentration was resuspended in cold, sterile PBS, aliquoted and stored at −80°C until required.

Prior to transduction, lentiviral particles were titrated using the qPCR lentivirus titration kit (Applied Biological Materials Inc.) according to manufacturer’s instructions. BMMs were transduced with lentiviral particles in the presence of 20µg/ml protamine sulfate (Sigma-Aldrich) 24h after the first RANKL (10ng/ml, R&D Systems) stimulation of BMMs.

### Bone resorption and extracellular acidification assays

For bone resorption assays, primary BMMs were cultured directly on devitalized bovine cortical bone slices (Boneslices.com) for 10-12 days. Osteoclasts were visualized by incubating with TRAP stain at 37°C for 30min. Cells were then washed with PBS, and imaged using a standard, upright light microscope. To expose the underlying resorption pits, cells were mechanically removed by gentle brushing of the bone surface with a wet cotton bud. Slices were then washed with 70% ethanol, rinsed in PBS, and left to air-dry overnight. For detection of resorption pits using light microscopy, surfaces of the bone slices were stained with 1% toluidine blue for 30sec, excess toluidine blue removed by blotting with tissue paper and then imaged using the Nikon Eclipse T200 reflective microscope (NIKON) at 10x magnification. Images were captured in JPG format using Nikon digital sight DS-5MC and NIS BR Elements 3.2 software. Qualitative analysis of the resorption areas was performed using ImageJ (Fiji) software. To detect resorption pits using SEM, resorbed bone slices devoid of cells were mounted onto aluminium studs and imaged using the TM4000 Plus tabletop SEM (Hitachi). For confocal microscopy, cleaned bone slices were incubated in 0.2% BSA-PBS containing rabbit anti-collagen I antibody (1/300, Abcam) for 2h at RT. Slices were washed in PBS and subsequently incubated with Alexa Fluor 568 goat anti-rabbit IgG (1/500, Thermo Fisher Scientific) and IVISense Osteo 680 fluorescent probe (Perkin Elmer) for 1h at RT. Bone slices were washed in PBS before mounting onto glass slides with ProLong^TM^ Diamond antifade mountant and left to cure overnight. The samples were imaged using 100x oil immersion lenses. Image stacks were opened in Fiji, and maximally projected after using Temporal-Color Code to differentially pseudo-color each z depth from the start to the end of the image stack. For the assessment of extracellular acidification activity, osteoclasts were differentiated on bone-mimicking substrates (Osteo Assay Stripwell microplate, Corning) for 7 days after which Von Kossa staining was performed to visualize the remaining mineralized matrix. Briefly, osteoclasts were removed by incubating in a 25% bleach solution in distilled water for 5min at RT. Wells were rinsed twice with distilled water before 100µl 1.5% silver nitrate (Sigma-Aldrich) was added to each well and incubated in the dark for 5min at RT. The silver nitrate solution was removed and the wells were rinsed well with several changes of distilled water. An equal volume of freshly prepared 0.5% hydroquinone was then added to each well. Precipitation reaction resulted in the remnant mineral turning dark brown/black upon addition of hydroquinone. After a 1min incubation, the hydroquinone solution was removed and the wells were washed gently with distilled water. Demineralized zones appear as white clearings against black mineral background and correlate with the degree of extracellular acidification by osteoclasts.

### Histology and histomorphometry

Left proximal tibia and distal femur samples from sex and age matched wild-type and knockout littermates were processed using a Leica TP1020 processor (Leica Biosystems) in preparation for methyl methacrylate (MMA) embedment according to standard protocol. Samples were sectioned using a Leica Biosystems RM2255 automated microtome at a thickness of 5µm. Sections were stained for TRAP, Mason’s trichrome, Von Kossa, according to standard protocols. Sections were scanned using a Leica Biosystems Aperio ScanScope. Histomorphometric analysis was performed using BioQuant Osteo, version 13.2.6 (Bioquant Image Analysis, Nashville, TN). For dynamic bone histomorphometry, 11-week-old mice were injected (intraperitoneally) with 5mg/kg fluorochrome-labeled calcein green (Sigma-Aldrich) twice over 6-day interval (i.e. on day 0 and then day 5). Mice were sacrificed 2 days after the second injection. Femora and tibia bone tissue were removed, fixed, embedded in SCEM-L1, cryosectioned and slides imaged by confocal microscopy. The mineral apposition rate (MAR) in micrometers per day were calculated from fluorochrome double calcein labels at the periosteal surfaces using NIS Elements software (Nikon).

### Micro-computed tomography (μCT)

Femurs, tibia, vertebrae and skulls were scanned at a voxel size of 8.89µm using Skyscan1176 (Bruker, Kontch, Belgium) Version1.1 (build 10) at X-ray source voltage 50kVp and current 500µA. Scans were reconstructed with NRecon version 1.6.10.4 (64bit) then analyzed with Bruker CTAn (CT-analyser) version 1.14.4.1. 300 slices below the growth plate were analyzed for the trabecular bone and 100 slices for the cortex. 3D models of tibias and femurs were visualized using Skyscan CTvox software version3.0.0r1114 (Bruker, Kontich, Belgium).

Analysis were performed on trabecular bone volume, trabecular thickness, trabecular separation and trabecular number. Cortical bone analysis was performed on cortical thickness.

High resolution µCT was performed on 12-week-old femurs to assess periosteal bone lesions using a Carl Zeiss X-ray Microscope Versa 520. In this instance, bone samples were scanned at 3µm resolution using Versa 520 v10.6.2005.12038, then reconstructed with XMReconstructor v10.7.36.79.13921 and finally 3D models assembled with TXM3DViewer version.

### Three-point bending test

Femurs were collected from 12-week-old WT and *Slc37a2*^KO^ male mice in saline solution and stored at −80°C. Bones were slowly thawed at 4°C and then warmed to room temperature. Femur lengths and diameters were measured using callipers. The three-point bending test was performed using an Instron 5566 testing machine (Instron Pty Ltd) with a 100N load cell and a custom designed bending rig with a 10mm span at the midshaft of femurs with a displacement rate of 1mm/min until the bone fractured with the force and displacement acquired digitally. The ultimate load (N) and stiffness (slope of the linear part of the curve, representing the elastic deformation in N / mm) of the midshaft were calculated according to the corresponding load and femur diameter.

### Metabolite extraction

Plates containing cells were thawed on ice. For metabolite extraction, 400µl of ice cold 9:1 of methanol:chloroform mixture (containing 4µl of 1mM D-Sorbitol-13C6 and 4µl 1mM L-Valine-13C5,15N as internal standard) was added, followed by rapid scraping of cells using a cell scraper (Sarstedt) and collected into 2ml Eppendorf tubes. 100µl of chloroform (Sigma Aldrich) was added immediately vortexed vigorously for 15sec before shaking on an ice block for 15min. Samples were then centrifuged at 16,000 x g at 4°C for 3min. The polar phase was collected into fresh tubes and 150µl of water (GC-MC grade, Sigma Aldrich) was added to each samples and vortex at high speed for 1min. Samples were centrifuged and 300µl of aqueous phase collected into GC-MS glass vials to dry in a SpeedVac (Labconco) at RT overnight. Data acquisition of metabolite profiles was performed as described by (Dias et al., 2015) with slight modifications. Polar metabolite extracts were derivatized by methoxyamination and trimethylsilylation using a Gerstel MPS2 XL automated sampling system. During this process the samples were shaken for 2h at 37 °C and 650rpm in 20μl of MeOX before being incubated for 1h at 37 °C and 650rpm in 30μl of methyl-trimethylsilyl trifluoroacetamide (MSTFA) mixture.

Metabolite profiles were acquired utilizing a 7890A Gas Chromatograph coupled with a 5975C Mass Selective Detector (Agilent Technologies, Santa Clara, CA, USA). 1µl of sample was injected onto a Agilent VF-5msec (J&W, Australia) in splitless mode and inlet temperature of 280°C. Helium was used as the carrier gas at a flow rate of 1ml/min. Oven conditions were set at 70°C starting temperature, held for 1min, then ramped at 7°C/min to 325 °C and held for 5min. Data acquisition was controlled by the Agilent MassHunter software package. Chromatogram processing and metabolite identification was performed utilizing AMDIS software (v 2.73), the in-house Metabolomics Australia mass spectral library and NIST14 database (National Institute of Standards and Technology, Gaithersburg, MD, USA). The putative results were normalized by the abundance of internal standard in each sample and corrected for weight.

### Electron microscopy

Transmission electron microscopy was carried out essentially as described by the methods of (Pavlos et al., 2005). Briefly, long bones from 5-day-old WT and *Slc37a2*^KO^ littermates were dissected free of soft tissue and fixed with 2.5% glutaraldehyde, 4% paraformaldehyde in 0.01 M cacodylate buffer (pH 7.3) for at least 1h at room temperature. After removal of the fixation solution, bones were decalcified for 5-days in 14% EDTA (pH 7.4) containing 0.1% glutaraldehyde. Subsequently, bones were washed, cut into ∼ 1 mm slices just below the growth plate and postfixed with 1% osmium tetroxide, dehydrated in ethanol, and embedded in Epon 812 (Nissin EM, Tokyo). Ultrathin sections were counterstained with lead citrate and examined with a Philips 410 LS transmission electron microscope (Phillips Inc, Eindhoven, The Netherlands). For assessment of SPION-enriched SL fractions, 6µl of the F2 fraction was absorbed onto Formvar-coated 150 mesh copper grids (ProSciTech) and imaged as described above.

### Statistics

Statistical analysis was performed using GraphPad Prism (GraphPad Software 9.0.2, San Diego, CA). For the experimental cohorts, 3-16 mice per genotype per experiment were used (=biological replicates) as indicated in the figures/figure legends for each experiment. For mouse experiments, age and sex-matched wildtype littermates were used as controls. All experiments were performed at least 3 times with the indicated number of replicates. Sample sizes were based on experience and experimental complexity but no methods were used to determine normal distribution of the samples. Tests between two groups were carried out using unpaired, two-tailed Student’s t-test. Multiple comparisons were assessed using nested one-way ANOVA with Tukey’s posthoc analysis. Data are presented as mean ± standard deviation (SD). Significance was labeled with * = *P* < 0.05, ** *P* < 0.01, *** *P* < 0.001, and **** *P* < 0.0001.

## Data availability

The data reported in this paper is available from the corresponding author upon reasonable request.

