## Supplementary Information for "Sugar transporter Slc37a2 regulates bone metabolism via a dynamic tubular lysosomal network in osteoclasts"

**EXTENDED DATA**

### EXTENDED DATA METHODS

#### Cell Culture

Immortalized human embryonic kidney cells (HEK293) and baby hamster kidney (BHK) cells were passaged in 10% FBS in Dulbecco's modified eagle medium (Life Technologies) supplemented with 2mM L-glutamine (Life Technologies). Cells were transfected with Lipofectamine 2000 (Life Technologies).

#### Skeletal staining

Sacrificed P5 mice were skinned, eviscerated, and fixed in 90% ethanol for 7 days and prepared for whole mount staining with Alcian blue and Alizarin red S according to the protocol detailed in (Chan et al., 2016).

#### RNAseq

For RNAseq analyses, femurs and tibias were dissected from three male and female WT and *Slc37a2*$^{KO}$ mice followed by whole-bone RNA extraction. Briefly, bones were frozen in liquid nitrogen and crushed before the addition of Trizol with vigorous shaking. Chloroform was added, and the samples were spun down at 12,000rpm for 30min at 4°C. The aqueous phase was collected and mixed with one volume of 70% ethanol. Subsequent RNA extraction steps were performed with the Life Technologies RNA extraction kit (Cat No. 12183018A). RNA sequencing was performed through the Illumina sequencing service by the Australian Genome Research Facility. Image analysis was performed in real time by the NovaSeq Control Software (NCS) v1.7.0 and Real Time Analysis (RTA) v3.4.4, running on the instrument computer. RTA performs real-time base calling on the NovaSeq instrument computer. Illumina bcl2fastq 2.20.0.422 pipeline was used to generate the sequence data. The primary bioinformatics analysis involved demultiplexing and quality control (QC). The data was then processed through an RNA-seq expression analysis workflow, which included alignment, transcript assembly, quantification, and normalisation. Analysis of the Top 1000 differentially expressed genes was performed.

#### Confocal microscopy

HEK293 cells expressing $^{emGFP-}$Slc37a2 were fixed and immunostained with antibodies against CD63/LAMP3 (monoclonal mouse DSHB Cat#. H5C6, 1/100 dilution) and GM130 (monoclonal mouse Clone 35/GM130 (ROU) BD Biosciences, CAT#. 610822, 1/100 dilution according to the methods described above. In some instances, $^{emGFP-}$Slc37a2 expressing cells were labeled with LysoTracker™ Red DND-99 (Thermo Fisher Scientific) probes or co-transfected with late-endosomal/lysosomal markers ($^{\text{mRFP-}}$Rab7 or LAMP1$^{\text{-RFP}}$) and imaged live using the NIKON A1R in a Ti-E inverted motorized confocal under controlled atmospheric conditions (37°C and 5% $CO_2$) using an Oko Labs stage top incubator. Frames were captured at 8sec intervals using NIS Elements software (Nikon). Images were processed and channels pseudo-colored using ImageJ (Fiji) software (Schindelin et al., 2012).

**APEX Electron microscopy**

Ascorbate peroxidase (APEX) processing of BHK cells co-transfected with $^{\text{emGFP-}}$Slc37a2 and APEX-GBP (GFP-binding peptide) was performed according to the methods described in (Ariotti et al., 2015). Briefly, transfected cells were fixed in 2.5% glutaraldehyde, washed repeatedly in 0.1M sodium cacodylate buffer prior to the DAB reaction. Cells were incubated with DAB in the presence of $H_2O_2$ for 30min at RT, washed in 0.1 M sodium cacodylate buffer, post-fixed in 1% osmium tetroxide for 2min, washed again, then serially dehydrated in increasing percentages of ethanol. Cells were serially infiltrated with LX112 resin in a Pelco Biowave microwave then polymerized overnight at 60°C. Ultrathin sections were cut on an ultramicrotome (Leica EM UC6, Leica Microsystems) and imaged using a JEOL1011 electron microscope (JEOL) at 80kV.

### EXTENDED DATA FIGURE LEGENDS

**Extended Data Fig.1 Slc37a2$^{+}$ tubular SLs colocalize with endolysosomal markers in living osteoclasts**

(**A**) Representative confocal images of live BMM-derived mouse osteoclasts on glass expressing $^{emGFP-}$Slc37a2 isoform 2 together with indicated organelle markers. Arrows denote co-labeled organelles in magnified enlargements. Bar, 10 µm.

(**B**) Colocalization analyses as determined by Pearson's correlation coefficient (Rr) (*n*= 8-12 cells per group).

**See also Videos S1 & S2**

**Extended Data Fig.2 Tubular SLs in naïve mouse osteoclasts. Related to Fig. 2.**

(**A**) Confocal images of a live mouse BMM-derived osteoclast culture on glass and probed with LysoTracker Green and DQ-BSA.

(**B**) Confocal microscopy images of live mature primary osteoclast grown on glass pulsed with LysoTracker Red and SIR-actin with enlarged time-lapse inlay.

(**C**) Electron micrograph of a representative mouse osteoclast lining trabecular bone within the primary spongiosa of 5-day-old mice. Magnified picture illustrate tubular SL-like structures (green arrows) in the vicinity of the nascent ruffled border.

**See also Video S5**

**Extended Data Fig.3 Generation and phenotyping of *Slc37a2*$^{KO}$ mice. Related to Fig. 4.**

(**A**) Schematic of the *Slc37a2*$^{tm2a(KOMP)Wtsi}$ knockout first allele targeting strategy.

(**B**) Genotyping confirming disruption of the *Slc37a2* gene locus. WT, wildtype (+/+); HET, heterozygous (-/+); KO, homozygous (-/-).

(**C**) qPCR confirmation of *Slc37a2* mRNA depletion in femurs (*n* = 3).

(**D**) Body weight of age and sex matched WT and *Slc37a2*$^{KO}$ mice. Male mice (WT, *n* = 8-18 per time point) and female mice (WT, *n* = 5-9 per time point).

(**E**) Five-day-old whole-mount skeletal preparations stained for Alcian blue (cartilage) and Alizarin red (bone).

(**F**) Whole body mammography of 12-week old female *Slc37a2* WT, HET and KO mice.

(**G-H**) Representative µCT reconstructed images of femurs (**G**), skulls (**H**) from 12-week-old female mice with genotypes indicated.

(**I**) Quantification of femur length of 12-week-old-female *Slc37a2* WT (*n* = 7), HET (*n* = 22) and KO (*n* = 10) mice.

(**J-N**) Representative sagittal µCT section (**J**) and µCT analysis (**K-N**) of distal femurs of 12-wk-old male (*n* = 8-10) and female (*n* = 6-9) *Slc37a2* WT, HET and KO mice.

(**O**) Macroscopic overview of organs from WT and *Slc37a2*$^{KO}$ mice.

(**P**) Representative H&E staining of paraffin-embedded tissue sections of WT and Slc37a2$^{KO}$ mice. Original magnification: ×100.

All data represent mean ± SD. NS=not significant ** $P < 0.01$, *** $P < 0.001$ by Student's t test (**C, D**) and by one-way ANOVA (**I, K-N**).

**Extended Data Fig.4 *In vitro* osteoblast differentiation and function is unaltered in *Slc37a2*$^{KO}$ mice. Related to Fig. 5.**

(**A**) Representative ALP and alizarin red staining of WT and *Slc37a2*$^{KO}$ osteoblasts isolated from calvarial bone following culture in osteogenic media (50 µg/ml L-ascorbic acid, 2 mM β-glycerophosphate, $10^{-8}$ M dexamethasone) for 21-days.

(**B**) Alizarin red staining of WT and *Slc37a2*$^{KO}$ of mesenchymal stem cell-derived osteoblasts isolated from long bones cultured *in vitro* in the absence or presence of osteogenic media for 21 days.

**Extended Data Fig.5 Multi-omic analyses of wild-type and *Slc37a2*$^{KO}$ osteoclasts. Related to Fig. 6.**

(**A**) Schematic of the sample acquisition and omic tools used to compose the osteoclast proteome, transcriptome and metabolome. Created with BioRender.com.

(**B**) Volcano plot summarizing the differences between the WT and *Slc37a2*$^{KO}$ osteoclast transcriptomes (*n* = 3 per genotype).

(**C**) Magnification of the volcano plot in (**B**) summarizing the differences between the WT and *Slc37a2*$^{KO}$ osteoclast transcriptomes.

(**D**) Volcano plot summarizing the differences between the WT and *Slc37a2*$^{KO}$ osteoclast cellular proteomes (*n* = 3 per genotype).

(**E**) Volcano plot summarizing the differences between the WT and *Slc37a2*$^{KO}$ osteoclast SL proteomes (*n* = 3 per genotype).

(**F**) Venn diagram summarizing the intersection between up-regulated hits ($P$ >0.05, FC>1.5) from the transcriptomic and proteomic analyses;

(**G**) Venn diagram summarizing the intersection between down-regulated hits ($P$ >0.05, FC-<1.5) from the transcriptomic and proteomic analyses.

**Extended Data Fig.6 $^{emGFP-}$Slc37a2 localizes to late-endosomes/lysosomes in non-osteoclastic cell lines. Related to Fig. 2.**

(**A**) Representative confocal images of HEK293 cells expressing $^{emGFP-}$Slc37a2 with indicated organelle markers. Bar, 10 µm.

(**B-C**) Live cell confocal imaging of a HEK293 cell co-expressing $^{emGFP-}$Slc37a2 with LAMP1$^{-RFP.}$ Magnified region highlights colocalization of puncta and blue boxed region corresponds with time-lapse series shown for an individual endolysosome shown in (**C**) with associated intensity signal of fluorophores shown over time, intervals = 8 sec.

(**D**) High-resolution time-lapse series of an $^{emGFP-}$Slc37a2 labeled endolysosome co-labeled with Lysotracker Red. Corresponding line scans are shown below, intervals = 8 sec. Bar, 1 µm.

(**E**) Detection of $^{emGFP-}$Slc37a2 using a co-expressed APEX-tagged GFP-binding protein (APEX-GBP) in BHK cells results in prominent DAB reaction product labeling of late endosomes and lysosomes. Bar, 1 µm.

### TABLES

**Extended Data Table 1, The osteoclast secretory lysosome (SL) proteome. Related to Fig. 1.**

### SUPPLEMENTAL VIDEOS

**Video S1. Slc37a2 localizes to a dynamic network of acidified tubular lysosomes in osteoclasts which house cathepsin K, Related to Fig. 2.** Representative time-lapse video of $^{\text{emGFP-}}$Slc37a2 in osteoclasts with indicated marker proteins cortactin$^{\text{-dtTomato}}$ (**A**), LysoTracker (**B**), $^{\text{mCherry-}}$VAMP7 (**C**) and Cathepsin K Magic Red (MR) (**D**). Yellow circles denote nuclei.

**Video S2 .$^{\text{emGFP-}}$Slc37a2+ tubular SLs are morphologically and dynamically distinct from endosomes, Related to Fig. 2.** Representative time-lapse video of $^{\text{emGFP-}}$Slc37a2 and early endosome marker $^{\text{mCherry-}}$Rab5 in osteoclasts.

**Video S3. $^{\text{emGFP-}}$Slc37a2$^{+}$ tubular SLs are acidified and fuse with the osteoclast plasma membrane, Related to Fig. 2.** Representative time-lapse video of a $^{\text{emGFP-}}$Slc37a2 expressing osteoclast cultured on glass labeled with indicated markers. Yellow dashed line denotes plasma membrane and red arrows highlight fusion events in magnified view.

**Video S4. Slc37a2$^{+}$ tubular SLs infiltrate the ruffled border and fuse with the bone-apposed plasma membrane, Related to Fig. 2.** Representative time-lapse video of $^{\text{emGFP-}}$Slc37a2 in an osteoclast cultured on bone co-labeled with LysoTracker and viewed at the level of the ruffled border (yellow dashed outline). Arrows on magnified images highlight a fusion event.

**Video S5. Osteoclasts possess dynamic tubular SLs which store cathepsins and form a network within F-actin rings, Related to Fig. 2.** Representative time-lapse video of mouse BMM-derived osteoclasts cultured on either glass (**A**) or bone (**B**) and labeled with the indicated vital markers. Arrows denote ruffled border (yellow) and F-actin ring/sealing zone (green), respectively.

**Video S6. Invertase-induced sucrosome resolution and tubulation is impaired in *Slc37a2*$^{\text{KO}}$ osteoclasts, Related to Fig. 7.** Representative time-lapse video of LysoTracker-labeled sucrosomes in WT and *Slc37a2*$^{\text{KO}}$ osteoclasts 1 h after the addition of invertase to induced sucrosome resolution and membrane tubulation. Arrows highlight tubulation events.

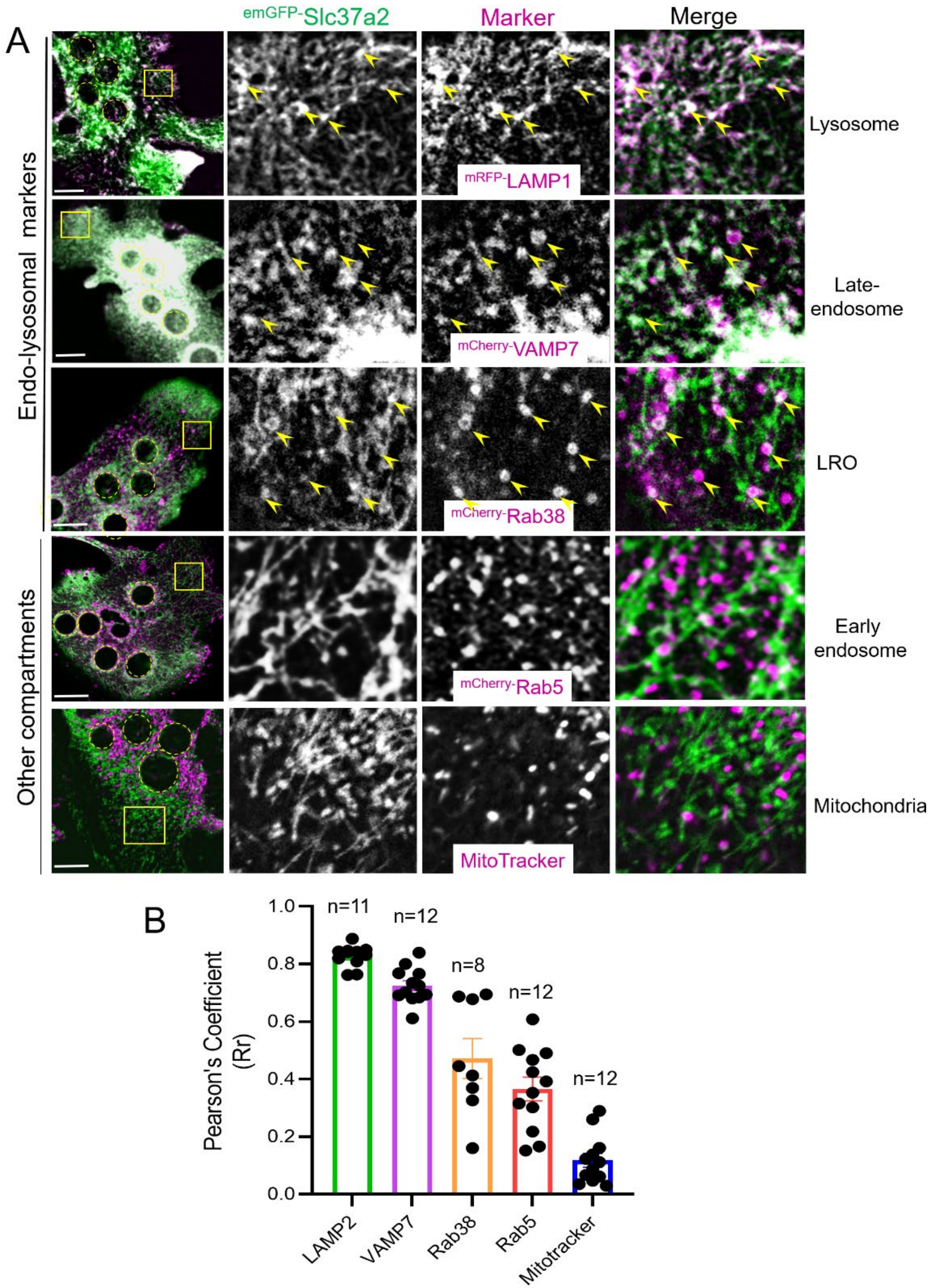


Extended Fig.1

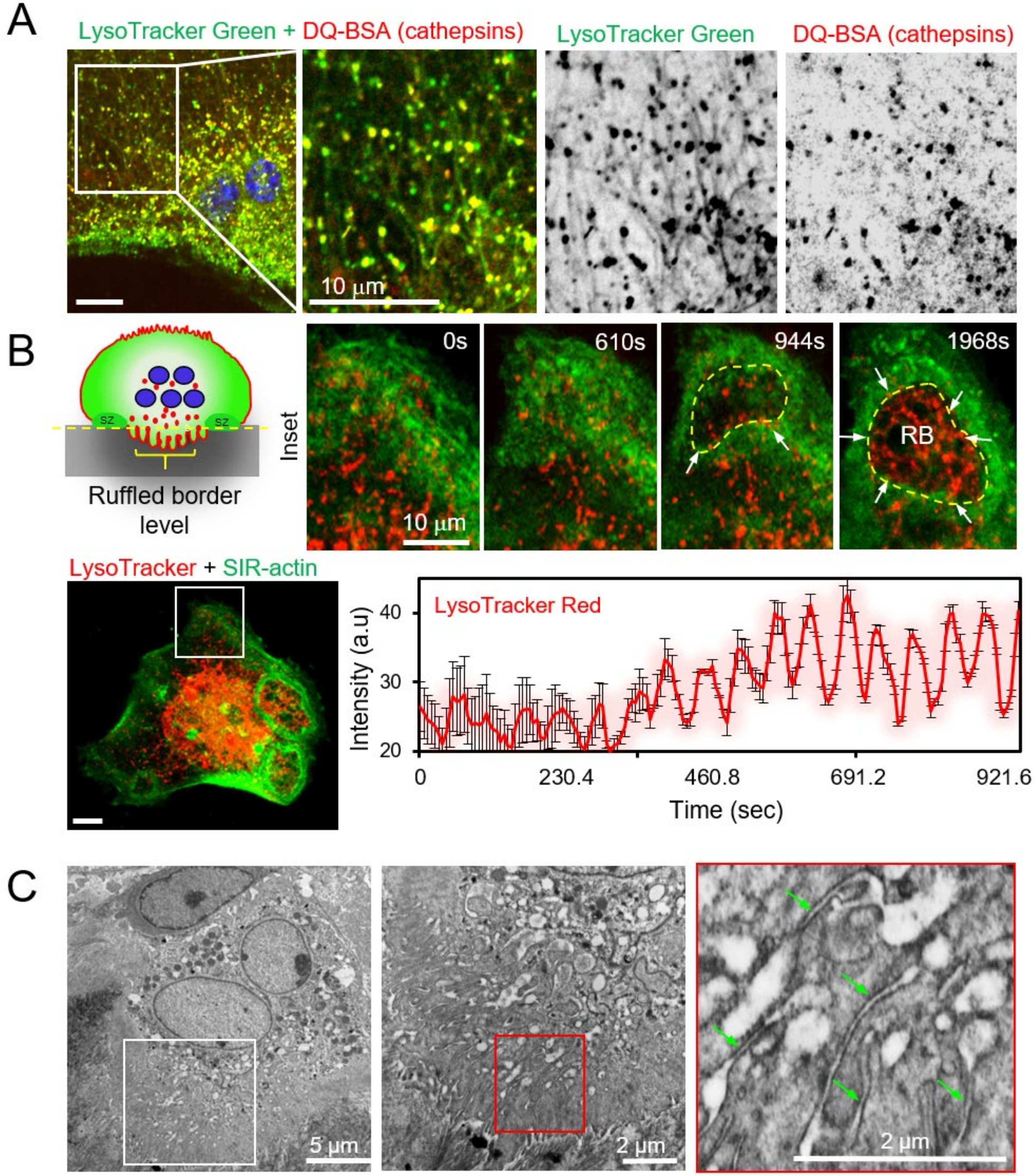


Extended Fig.2

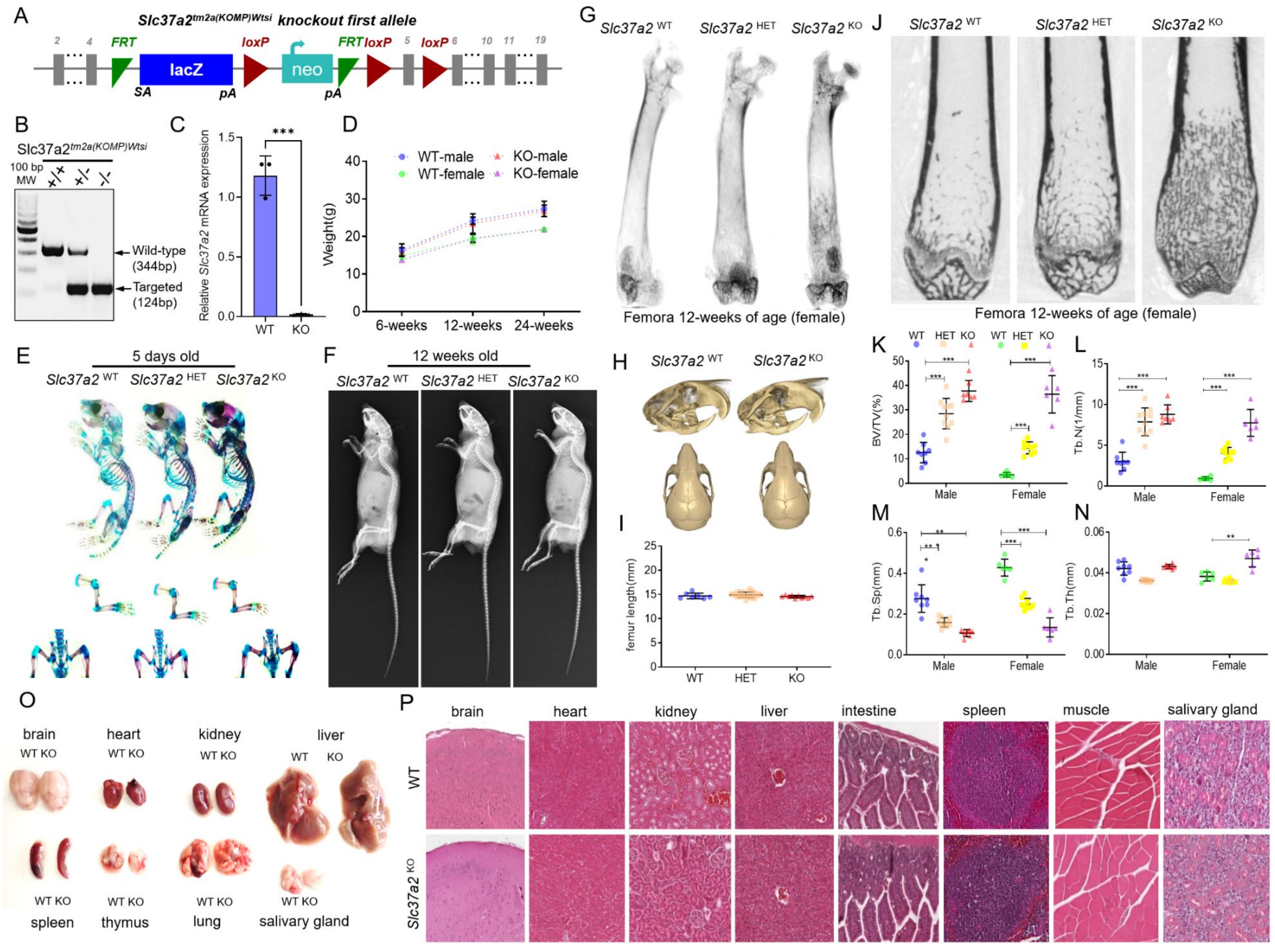


Extended Fig.3

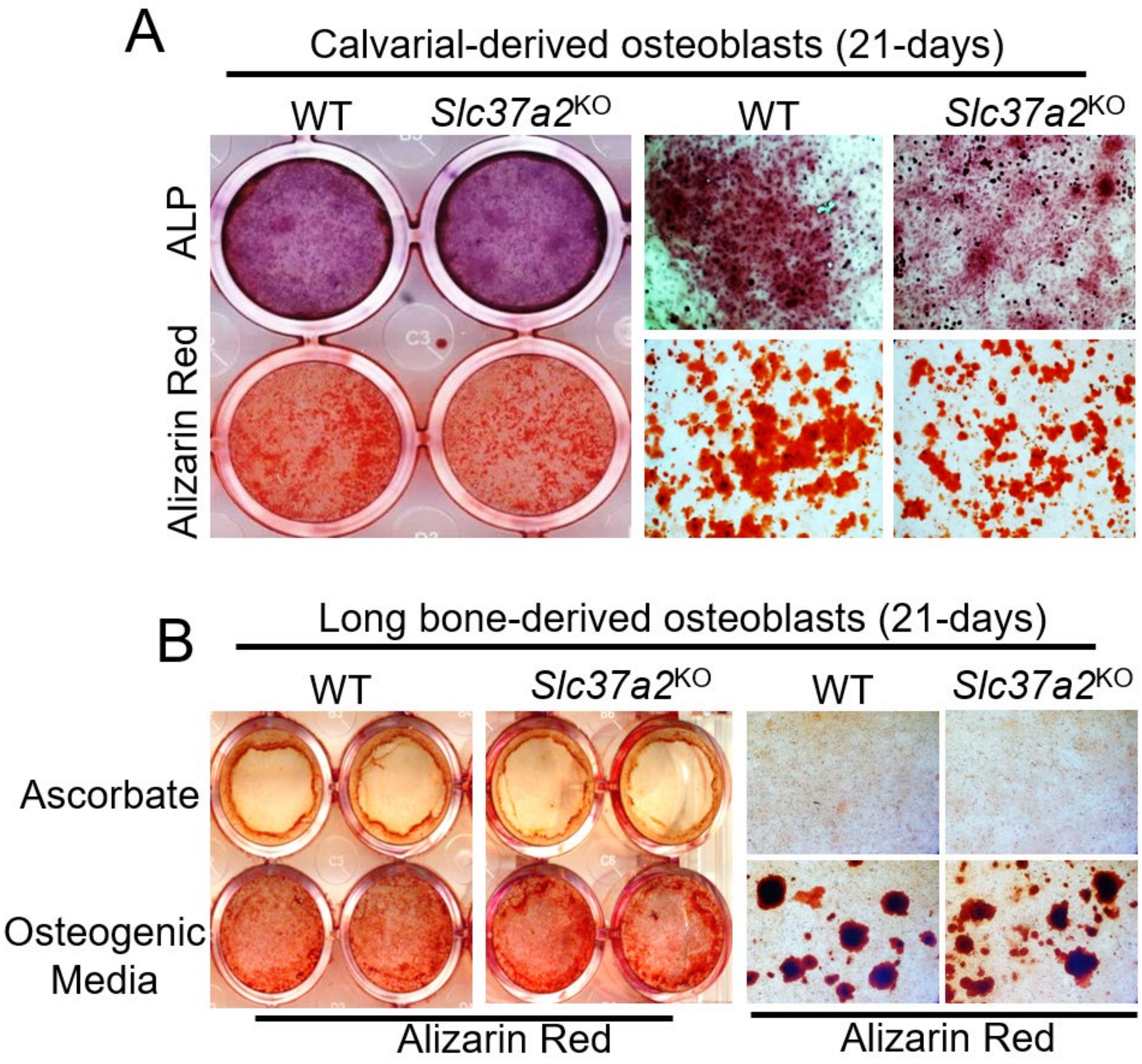


Extended Fig.4

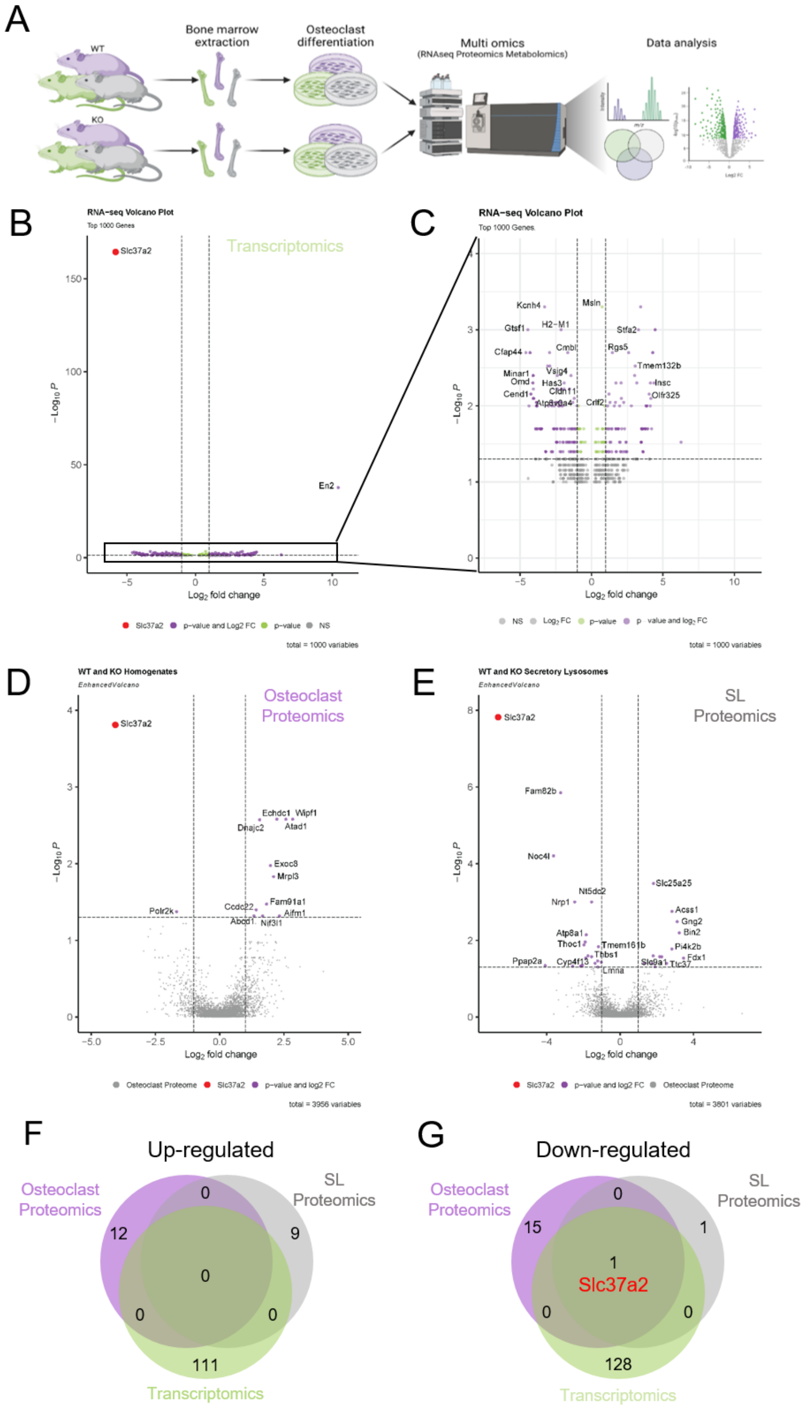


Extended Fig.5

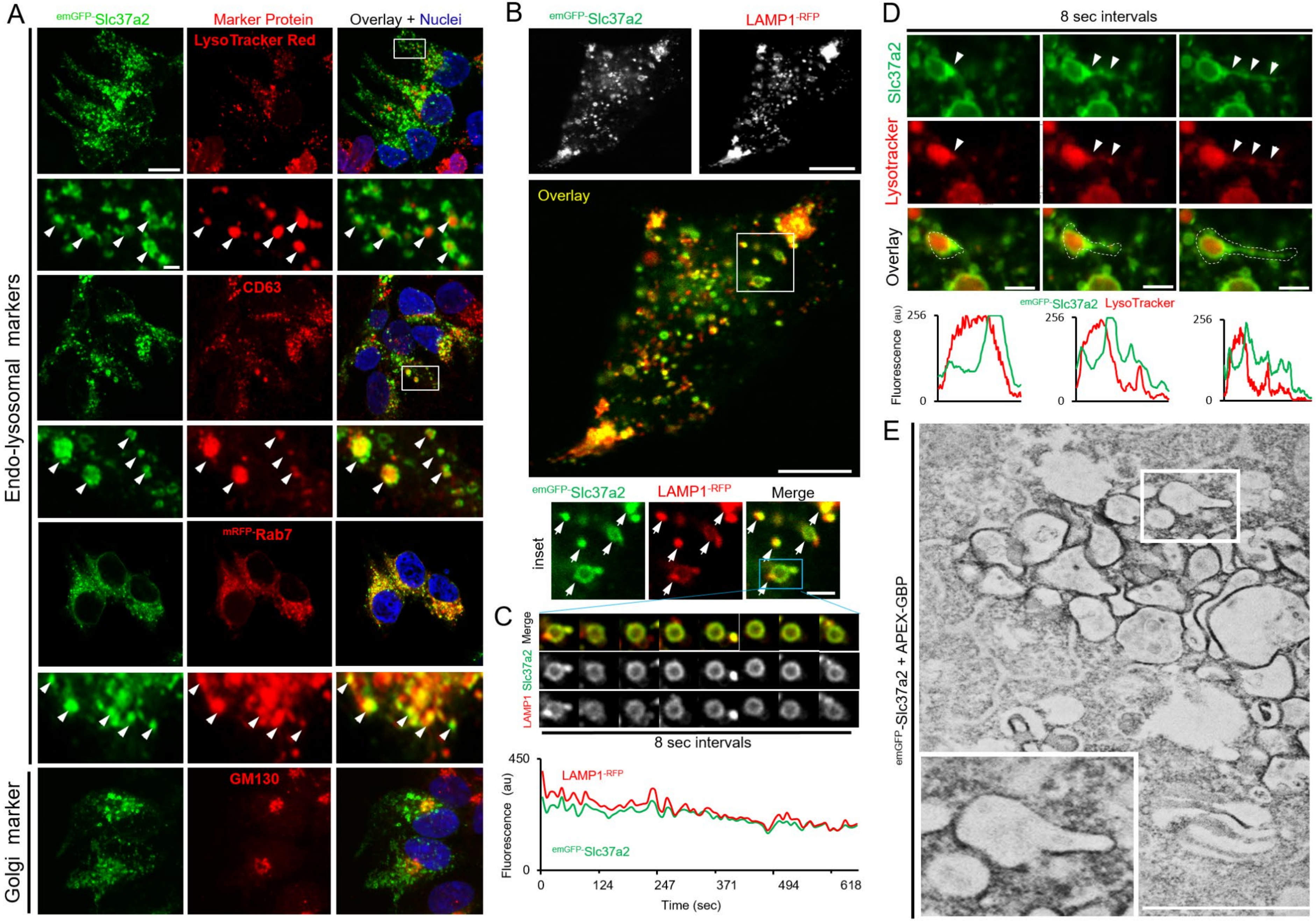


Extended Fig.6
